# Dorsal/NF-κB exhibits a dorsal-to-ventral mobility gradient in the Drosophila embryo

**DOI:** 10.1101/320754

**Authors:** Hadel Al Asafen, Natalie M. Clark, Etika Goyal, Sadia Siddika Dima, Hung-Yuan Chen, Thomas Jacobsen, Rosangela Sozzani, Gregory T. Reeves

## Abstract

Morphogen-mediated patterning is a highly dynamic developmental process. To obtain an accurate understanding of morphogen gradient formation and downstream gene expression, biophysical parameters such as protein mobilities must be quantified *in vivo*. The dorsal-ventral (DV) patterning of early *Drosophila* embryos by the NF-κB homolog Dorsal (Dl) is an excellent system for understanding morphogen gradient formation. Dl gradient formation is controlled by the inhibitor Cactus/IκB (Cact), which regulates the nuclear import and diffusion of Dl protein. However, quantitative measurements of Dl mobility and binding are currently lacking. Here, we use scanning fluorescence correlation spectroscopy to quantify the mobility of GFP-tagged Dl. We find that the DNA binding of Dl-GFP, which affects its mobility, varies along the DV axis, with highest DNA binding on the ventral side. Moreover, we also observe that the time scale for Dl-GFP to exit the nucleus is longer in the ventral and lateral regions of the embryo, which is consistent with stronger DNA binding. Using analysis of mutant alleles of *dl* tagged with GFP, we conclude that Dl-GFP/Cact interactions in the nuclei are responsible for the variation in Dl-GFP/DNA binding along the DV axis, which impacts our understanding of the spatial range of the Dl gradient and the robustness and precision of downstream gene expression. Thus, our results highlight the complexity of morphogen gradient dynamics and the ability of quantitative measurements of biophysical interactions to drive biological discovery.

## Introduction

Tissues in a developing organism are patterned by short and long-range signaling achieved through morphogen concentration gradients, which carry the positional information necessary to control gene expression in a spatially dependent fashion. In the past two decades, studies using GFP-tagged morphogens -- including early *Drosophila* morphogens Bicoid and Dorsal; Dpp in the wing imaginal disc; and the Hedgehog, Wnt, and TGF-β families in vertebrates -- have revealed that the establishment of morphogen gradients is a highly dynamic and complex process (Delotto et al., 2007; Entchev et al., 2000; Gregor et al., 2007; Holzer et al., 2012; Luz et al., 2014; Reeves et al., 2012; Ribes et al., 2010; Teleman & Cohen, 2000; Wallkamm et al., 2014; Wartlick et al., 2011; Williams et al., 2004; Zhou et al., 2012). To move toward a quantitative understanding of morphogen signaling and downstream gene expression patterns, several key biophysical parameters related to gradient formation, including diffusivity and binding, must be measured.

Dorsal (Dl), one of three *Drosophila* orthologs to mammalian NF-κB (Steward, 1987), acts as a morphogen to pattern the dorsal-ventral (DV) axis of blastoderm stage (1-3 h old) *Drosophila* embryos. Dl is initially distributed uniformly along the DV axis (Roth et al., 1989; Steward et al., 1988). After the 9^th^ nuclear division cycle, when the nuclei migrate to the periphery of the syncytial embryo, Dl begins to accumulate in ventral nuclei, and is excluded from dorsal nuclei (Roth et al., 1989; Steward et al., 1988). This DV gradient in Dl nuclear concentration is due to signaling through the Toll receptor on the ventral side of the embryo (Anderson, Bokla, et al., 1985; Anderson, Jürgens, et al., 1985; Belvin & Anderson, 1996). In the absence of Toll signaling, Dl remains bound by the cytoplasmic tethering protein Cactus (Cact), the *Drosophila* IκB homolog (Geisler et al., 1992; Kidd, 1992), and thus localized to the cytoplasm (Belvin et al., 1995; Bergmann et al., 1996; Reach et al., 1996; Whalen & Steward, 1993). On the ventral side, Toll signaling acts through Pelle kinase to phosphorylate Cact, which causes dissociation of the Dl/Cact complex and degradation of Cact (Daigneault et al., 2013). Once freed from binding to Cact, Dl enters the nucleus and regulates the expression of more than 50 target genes, which initiate further signaling pathways and specification of the embryo’s germ layers (Chopra & Levine, 2009; Moussian & Roth, 2005; Reeves & Stathopoulos, 2009; Schloop, Bandodkar, et al., 2020; Stathopoulos & Levine, 2002, 2004). Toll signaling also phosphorylates Dl, which has been shown to increase Dl nuclear localization (Drier et al., 1999, 2000; Gillespie & Wasserman, 1994; Whalen & Steward, 1993).

While this well-established mechanism rapidly initiates Dl gradient formation, recent work has revealed complex dynamics in the further maturation of the gradient. In particular, live imaging of fluorescently-tagged Dl proteins has shown that the Dl nuclear gradient grows slowly during interphase and collapses during mitosis (Delotto et al., 2007; Reeves et al., 2012). Quantification of the gradient levels has shown that over the course of nuclear cycle 10-14, Dl (both nuclear and cytoplasmic) accumulates on the ventral side of the embryo, causing the peak of the Dl gradient, in the ventral-most cells, to grow steadily over time (Carrell et al., 2017; Liberman et al., 2009; O’Connell & Reeves, 2015; Reeves et al., 2012; Schloop, Carrell-Noel, et al., 2020). In particular, diffusion and nuclear capture of Dl, both of which are regulated by Cact, are proposed to control the continuous growth of the Dl gradient peak (Carrell et al., 2017). Nonetheless, the diffusivity and nuclear transport rates are not quantitatively known. Furthermore, our previous work has suggested that Dl/Cact complex also is present in the nucleus, which has implications for the apparent spatial range of the Dl gradient, as well as for the robustness and precision of gene expression (Al Asafen et al., 2020; O’Connell & Reeves, 2015). Therefore, to fully understand the dynamics of Dl gradient formation and gene regulation, measurements of the biophysical parameters governing Dl mobility (specifically diffusivity, nuclear transport, and binding to DNA) are needed.

Here we employ scanning fluorescence correlation spectroscopy (scanning FCS) techniques to measure the mobility of Dl in live embryos. Scanning FCS techniques involve rapid and repeated acquisition of fluorescent imaging data using confocal microscopy. First, we apply a specific scanning FCS method known as Raster Image Correlation Spectroscopy (RICS) to autocorrelate GFP-tagged Dl over space and time in a small region of the embryo (Digman, Brown, et al., 2005; Digman, Sengupta, et al., 2005). The shape of this autocorrelation function is then analyzed to infer protein diffusion, which reveals two pools of Dl-GFP, mobile and immobile. Notably, we observed that the fraction of immobile Dl varies along the DV axis. Cross-correlation RICS (ccRICS), which uses two different fluorophores (GFP and RFP, in this case), suggests that the DV variation in immobile fraction is the result of a higher fraction of Dl binding to the DNA on the ventral side than on the dorsal side. In addition, we employ Pair Correlation Function (pCF) technique to compute the correlation between two pixels along a line scan, allowing us to visualize barriers to Dl movement (Digman & Gratton, 2009; Hinde et al., 2010). We show that the tendency of a Dl-GFP molecule to exit the nucleus also displays variation along the DV axis. When examining mutants that mimic the Dl gradient at different DV positions, the RICS and pCF data corroborate these findings. We propose two competing hypotheses to explain the DV variation in Dl/DNA interactions: that Toll phosphorylation of Dl may increase Dl/DNA interactions on the ventral side, or that Cact may bind Dl in the nuclei, which would prevent Dl from binding DNA in dorsal regions of the embryo. We test these competing hypotheses using mutant alleles of *dl* and conclude that Dl/Cact binding in dorsal nuclei prevents DNA binding, suggesting that further imaging of Cact may be needed to fully understand Dl-dependent gene expression. This result reveals a Dl gradient that has greater dynamic range, spatial range, robustness, and precision than previously understood (Al Asafen et al., 2020; Liberman et al., 2009; O’Connell & Reeves, 2015; Reeves et al., 2012; Schloop et al., 2025).

## Results

### Quantification of Dl Mobility Reveals a Dorsal-to-Ventral (DV) Gradient

To measure the mobility of Dl in the embryo, we performed Raster Image Correlation Spectroscopy (RICS) analysis on embryos carrying a monomeric GFP-tagged Dl (Carrell et al., 2017) and an H2Av-RFP construct in a *dl* heterozygous background (hereafter referred to as Dl-GFP embryos; see Materials and Methods). We performed RICS analysis in different locations along the DV axis (Fig. 1A) of nuclear cycle 12-14 embryos, when the Dl gradient is clearly defined and detectable. Specifically, we rapidly imaged small (pixel size ≤ 0.1 µm, 256×256 pixels; see Methods) regions of the embryo (Fig. 1B, Fig. S1A-D and Movies S1-3) and calculated 2D autocorrelation functions (ACFs; Fig. 1B and Equation 1 in Materials and Methods). The ACFs each have a fast (ξ) and slow (η) direction due to the rapid movement and line retracing of the confocal scan, respectively (Fig. 1C,D). We then fit a one-component, pure diffusion model to these ACFs (Fig. 1C and Equation 2 in Materials and Methods; see also Supplemental Information), which allowed us to estimate apparent diffusivities. We found that the apparent Dl diffusivity varied with the nuclear-to-cytoplasmic ratio (NCR) of Dl-GFP intensity, which we take as a proxy for position along the DV axis of the imaged region because embryos were mounted in random DV orientations (Fig. 1A). In particular, the apparent diffusivity was lower on the ventral side of the embryo than on the dorsal side (Fig. 1E). To test whether this trend is statistically significant, we used linear regression, and found the slope of the apparent diffusivity from ventral-to-dorsal to be statistically different from zero (p-value < 10^-4^; Fig. 1E).

**Figure 1:**
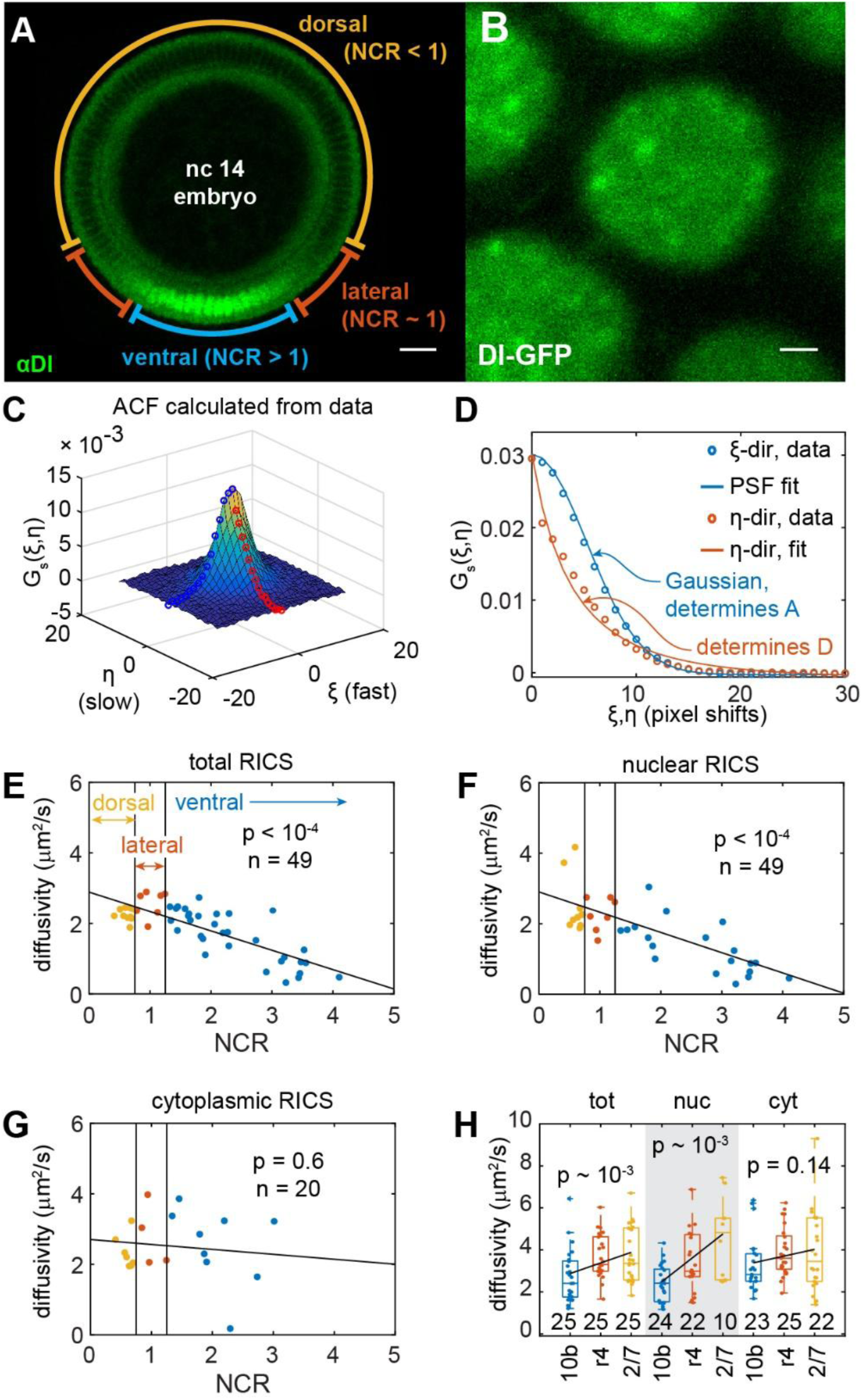
RICS analysis reveals a dorsal-ventral asymmetry in the mobility of Dl. (A) Representative image of the Dl gradient in an nc 14 embryo. Scale bar = 25 μm. (B) Representative image of Dl-GFP used for RICS analysis. Scale bar = 1 μm. (C,D) Plots of the autocorrelation function (ACF) of the image in A. (C) 3D plot of the ACF. Blue and red open circles represent the slice of the 3D surface for the fast and slow directions, respectively. (D) Plot of the fast and slow slices of the 3D ACF. Blue and red open circles (experimental data) correspond to those found in (C). The experimental data (open circles) are fit using a diffusion model (solid curves). The data and fit are separated into two components, the “fast” (ξ, blue) and “slow” (η, orange) directions. (E-G) Plots of the diffusivity of Dl-GFP vs NCR, measured using RICS on the entire imaging frame (nuclear plus cytoplasmic) (E), nuclear regions only (F), or cytoplasmic regions only (G). (H) Boxplot of the diffusivity of Dl-GFP in mutant embryos measured using RICS on the entire imaging frame (nuclear plus cytoplasmic) (“tot”; left side), nuclear regions only (“nuc”; center, gray), or cytoplasmic regions only (“cyt”; right side). In (E-H), blue represents ventral/“ventral-like”, orange represents lateral/“lateral-like”, and yellow represents dorsal/“dorsal-like” measurements. Solid dots represent individual measurements. Black lines represent a linear regression fitted to the data. *p*-values for the slope of the trendline being zero are indicated on the graph. Sample sizes indicated on graph.

Considering that, on the ventral side of the embryo, Dl is predominantly located in the nucleus, whereas on the dorsal side, Dl is predominantly cytoplasmic, it is possible that the spatial variation in the diffusivity arises from differences in the behavior of nuclear and cytoplasmic Dl. Therefore, we segmented the nuclei and calculated the ACF for the nuclei and the cytoplasm separately (see Supplemental Information). We found a ventral-to-dorsal trend in the apparent diffusivity of nuclear-localized Dl-GFP, as estimated by fitting to the pure diffusion model (p-value < 10^-4^, Fig. 1F). However, the apparent cytoplasmic diffusivities showed no statistically significant trend in spatial variation (p-value 0.6; Fig. 1G). These results suggest that the observed gradient in Dl mobility is specific to nuclear-localized Dl.

We reasoned that the gradient in the mobility of nuclear-localized Dl could be dependent on Toll signaling, which not only favors the dissociation of the Dl/Cact complex, but also phosphorylates Dl and increases its affinity for the nucleus (Drier et al., 1999, 2000; Gillespie & Wasserman, 1994; Whalen & Steward, 1993). To this end, we performed RICS analysis on three mutant lines with “ventral-like” (*Toll^10B^*; Schneider et al., 1991), “lateral-like” (*Toll^r4^*; Schneider et al., 1991), or “dorsal-like” (*pll^2/7^*; Anderson and Nüsslein-Volhard, 1984) levels of nuclear Dl-GFP (Fig. S1E-G). If the spatial gradient in Dl movement depends on Toll signaling, then we would expect *Toll^10B^* embryos to have the lowest apparent diffusivity, while *Toll^r4^* and *pll^2/7^* embryos would have higher apparent diffusivities. Accordingly, we found the apparent diffusivity of total (nuclear + cytoplasmic) Dl-GFP is correlated to the strength of Toll signaling across the mutants (Fig. 1H). Additionally, linear regression revealed a ventral-to-dorsal trend (p-value 10^-3^; Fig. 1H; left side). Similarly, the apparent diffusivity of nuclear Dl-GFP varies across these Toll signaling mutants with a statistically significant trend (p-value 10^-3^; Fig. 1H, center). In contrast, we did not see a variation in apparent cytoplasmic Dl-GFP diffusivity in the mutants (p-value 0.14; Fig. 1H, right side). It should be noted that these uniform Toll signaling mutants lack a shuttling mechanism that concentrates Dl on the ventral side of wildtype embryos (Carrell et al., 2017), and as such, the NCR observed in these mutants cannot be used as a direct positional variable for comparing mutants to wild type. For this reason, we presented the mutant results as boxplots to highlight overall trends rather than position-matched values. These results suggest that the variation of mobility along the DV axis is downstream of Toll signaling.

### The Gradient of Apparent Diffusivity is Due to Immobilized Dl in the Nucleus

Given that Dl is a transcription factor, it is likely that there are at least two pools of Dl: a freely diffusing pool, and a DNA-bound (immobile) pool. As such, it is possible that the diffusivity is constant, and the observed trend in apparent diffusivity, as inferred by the pure diffusion model, would in fact be a trend in the immobile fraction of Dl. For example, if a significant pool of immobile Dl exists, it would lower the apparent diffusivity, as estimated by a pure diffusion model.

To investigate this possibility, we fit a two-component diffusion model (Equation 3 in Materials and Methods) to the ACFs described above. In the two-component model, Dl could be in two states: freely diffusible and immobilized (Fig. 2A; see Supplemental Information for more details). Thus, an additional adjustable parameter is required: the fraction of immobilized Dl, ϕ. To avoid over-fitting, when using the two-component model, we fixed the diffusivity to be 3 μm^2^/s, consistent with the cytoplasmic diffusivity measured above (Fig. 1F, left side). In general, we found the two-component model fit the ACFs better than the one-component model (Fig. 2B). Furthermore, we found that *ϕ*_*nuc*_, the fraction of immobilized Dl in the nuclei, varies along the DV axis, with a statistically significant linear trend in NCR (p-value < 10^-4^; Fig. 2C). In contrast, in the cytoplasm, roughly 10-20% of Dl was immobilized (*ϕ*_*cyt*_) in wildtype embryos, and the variation in this fraction along the DV axis lacked a statistically significant trend (p-value 0.08; Fig. 2D).

**Figure 2:**
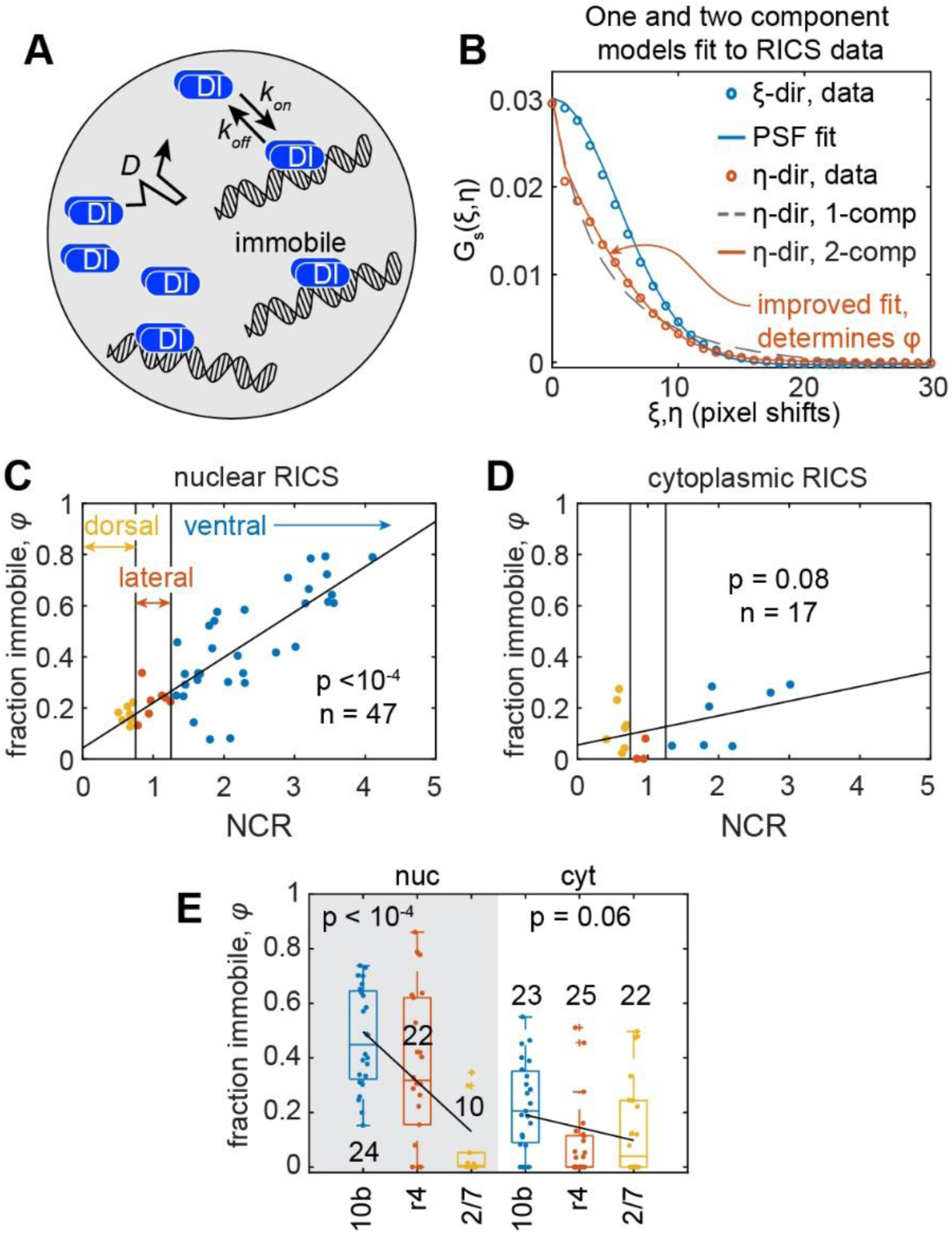
One- and two-component diffusion models quantify the relationship between DNA binding and mobility of Dl. (A) Illustration of the two-component model. Dl can diffuse and reversibly bind DNA. Dl bound to DNA is immobile. In contrast to the one-component diffusion model, in which Dl is freely diffusible and does not bind to DNA. (B) Comparison of one-component (dashed gray curve) and two-component (solid orange curve) diffusion models fit to the slow direction of the experimental RICS data (orange open circles). (C,D) Plots of the fraction of immobilized Dl (φ) vs. NCR measured using RICS on nuclear regions only (C) or cytoplasmic regions only (D). (E) Boxplot of φ in mutant embryos measured using RICS on nuclear regions only (“nuc”; left side, gray) or cytoplasmic regions only (“cyt”; right side). In (C-E), blue represents ventral/“ventral-like”, orange represents lateral/“lateral-like”, and yellow represents dorsal/“dorsal-like” measurements. Solid dots represent individual measurements. Black lines represent a linear regression fitted to the data. *p*-values for the slope of the trendline being zero are indicated on the graph. Sample sizes indicated on graph.

In the previously described Toll signaling mutants, the immobile fraction of nuclear Dl-GFP was statistically highly correlated with Toll signaling strength (p-value < 10^-4;^ Fig. 2E, left side). The immobile fraction in the cytoplasm in the Toll signaling mutants did not rise to a statistically significant trend (p-value 0.06; Fig. 2E, left side).

Taken together, our RICS data from both wildtype and mutant embryos are indicative of a two-component system where a pool of Dl is immobilized, and that the immobile pool, specifically in the nuclei, is highly correlated to Toll signaling. If the immobile pool represents Dl/DNA binding, then the expectation for a transcription factor interacting with DNA is that the concentration of immobile Dl should be a saturating, increasing function of total nuclear Dl concentration. In contrast, in such a saturating system, the *fraction* of immobile Dl, ϕ, is expected to decrease (or, at best, remain constant) with increasing nuclear total Dl concentration. However, we are observing the opposite trend (ϕ being positively correlated with nuclear Dl), suggesting that the variation in the immobile Dl pool along the DV axis is dictated by more than simply the nuclear Dl concentration.

### Dorsal/DNA binding exhibits a spatial gradient

Our RICS results suggest that the spatial variation in nuclear Dl movement is due to a population of immobilized Dl-GFP on the ventral side of the embryo. To test whether the immobilized Dl in the nuclei is due to DNA binding, and whether there is a spatial gradient in DNA binding, we used cross-correlation RICS (ccRICS; Digman et al., 2009) to measure the extent to which Dl (Dl-GFP) and Histone (H2Av-RFP) may be bound to the same physical structure (in this case, DNA). The ratio of the cross-correlation function (CCF) amplitude (Fig. 3A,B) to the ACF amplitude in the red channel (H2Av-RFP channel; Fig. 3C) is proportional to the fraction of Dl bound to DNA, ψ (see Supplemental Information)(Rippe, 2000; Weidemann et al., 2002). Note that it is possible for the CCF amplitude to be negative (Dl-GFP avoidance of H2Av-RFP), in which case, the fraction ψ could be viewed as zero. We found ψ has a statistically significant linear trend with NCR (p-value < 10^-4^; Fig. 3D). In some cases with high ψ, the overlap between Dl-GFP and H2Av-RFP can be seen visually (Fig. S2A and Movie S4). Furthermore, in dorsal regions, the correlation between histone and Dl is indistinguishable from zero. In these cases, the low correlation can be seen as the lack of peak in the CCF (Figs. 3B, S3B,C).

**Figure 3:**
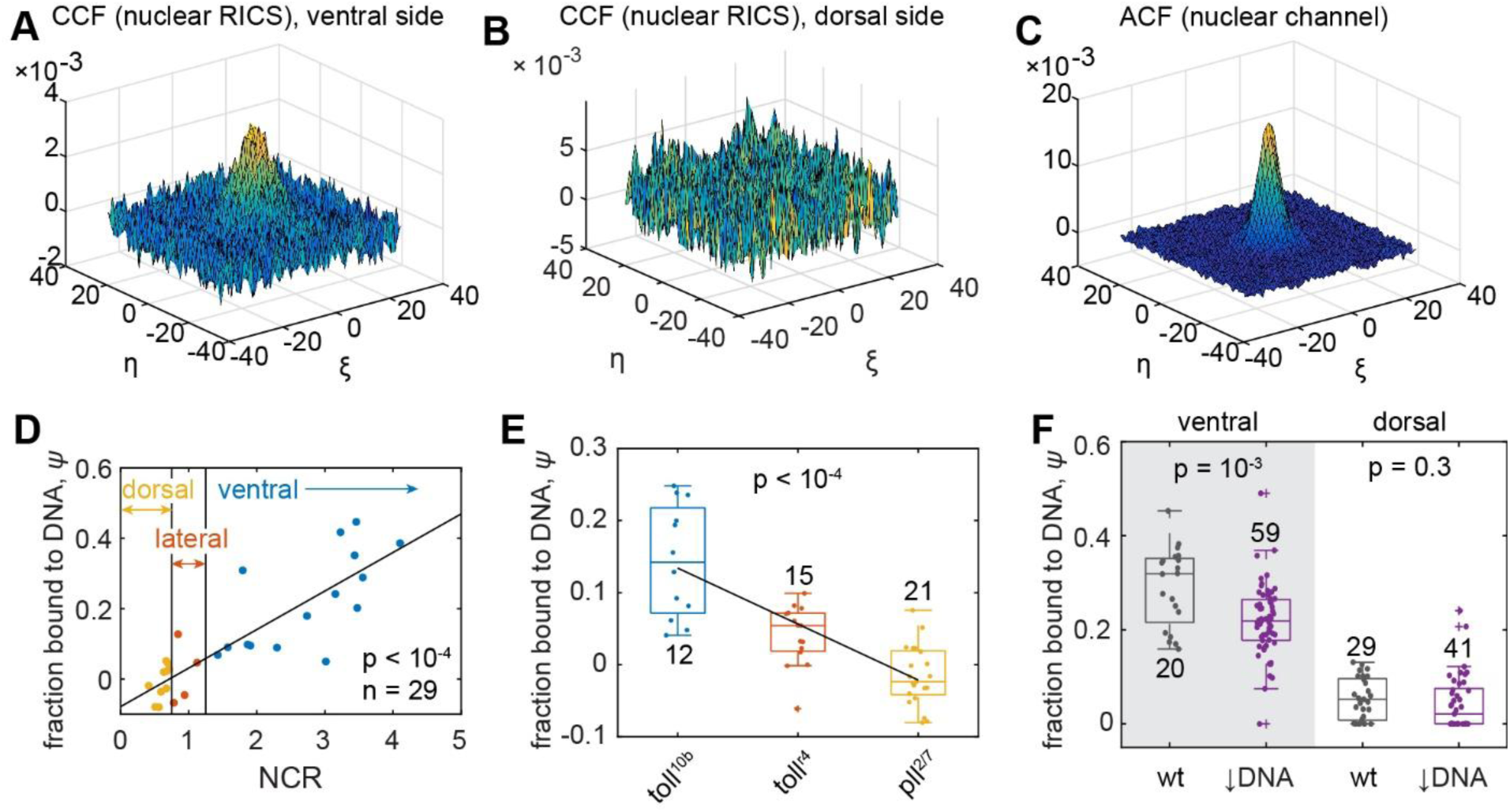
Cross-correlation RICS analysis of DNA-bound Dl. (A,B) Cross-correlation between Dl-GFP and H2Av-RFP, with nuclear mask, on the ventral side (A) or the dorsal side (B). (C) Example of autocorrelation function of H2Av-RFP, with nuclear mask. (D) Plot of ψ, the fraction of Dl bound to DNA, in wildtype embryos vs. NCR. (E) Boxplot of ψ in Toll signaling mutant embryos. In (D,E), blue represents ventral/“ventral-like”, orange represents lateral/“lateral-like”, and yellow represents dorsal/“dorsal-like” measurements. Solid dots represent individual measurements. Black lines represent a linear regression fitted to the data. *p*-values for the slope of the trendline being zero are indicated on the graph. (F) Boxplot of ψ, the fraction of Dl bound to DNA, in either wildtype (wt) embryos or embryos overexpressing a mutated form of Dl (R063C) with reduced DNA binding (↓DNA). Left side of plot (gray): ventral side; right side of plot (white): dorsal side. *p*-values for differences in means of the distributions are indicated on the graph. In (D-F): Sample sizes indicated on graph.

In the Toll signaling mutant embryos, ψ is also correlated with the levels of Toll signaling with a statistically significant trend (p-value < 10^-4^; Fig. 3E). Furthermore, ψ is indistinguishable from zero in *pll^2/7^* mutants. Note that, as with the Dl diffusivity, there may not be a direct comparison between ψ in the uniform Toll signaling mutants and the corresponding locations in wildtype embryos.

To test whether ccRICS is an appropriate assay to detect Dl/DNA binding, we made a GFP-fused allele of *dl* in which arginine 63 has been replaced by a cysteine (*dl*^R063C^-*gfp*), which is in the highly conserved DNA binding portion of the Rel homology domain and has been shown to severely reduce Dl-mediated gene expression (Govind et al., 1996; Isoda et al., 1992). As this allele is antimorphic (Isoda et al., 1992), we used a *nos*-Gal4 line to drive the expression of UAS*-dl*^R063C^-*gfp* (Brand & Perrimon, 1993) and overexpressed it in a maternal background of one copy of transgenic *dl-mGFP* and zero copies of endogenous *dl* (see Materials and Methods). To directly compare to a wildtype control, we also created a Gal4/UAS-driven wildtype allele of *dl* fused to *gfp* (UAS-*dl^wt^-gfp*). As predicted, the UAS-*dl*^R063C^-*gfp* allele had a statistically lower value of ψ on the ventral side than the wildtype control (p-value 10^-3^; Fig. 3F, left side). In contrast, both alleles had a low value of ψ on the dorsal side (p-value 0.3; Fig. 3F, right side). These results are consistent with ψ being a measure of Dl/DNA binding.

Overall, the findings that both the DNA-bound pool and the immobilized pool of nuclear Dl-GFP are correlated to both Toll signaling and the NCR suggest that DNA binding of Dl may be responsible for its overall lowered mobility on the ventral side of the embryo. Furthermore, as mentioned in the previous section, the trends of immobile fraction, ϕ, and fraction of Dl/DNA interactions, ψ, both being positively correlated with nuclear Dl concentration is the opposite of what would be expected for a saturating system consisting only of transcription factor/DNA interactions.

### Dl Exhibits DV Variation in Nuclear Export Propensity

Our RICS analysis revealed that Dl movement is lowest on the ventral side, where it is primarily nuclear localized and a significant fraction is immobile, and highest on the dorsal side, where it is primarily cytoplasmic. Thus, we reasoned that the ability of Dl to exit the nucleus could also vary spatially. To determine whether there is spatial variation in Dl movement into or out of the nucleus, we used Pair Correlation Function (pCF) analysis to measure any restriction of movement of Dl-GFP across the nuclear envelope (Digman & Gratton, 2009, 2011; Hinde et al., 2010). Here it is important to state the term "movement" can be ambiguous, and in other sections, movement refers to apparent diffusion, which is random, passive motion inside the nucleus or cytoplasm In this section, movement refers to the propensity of Dl to exit the nucleus.

In contrast to RICS, which uses 2D raster scans of the embryo, pCF requires line scans. For our purposes, these line scans cross one or more nuclei (white arrows in Fig. 4A, top row). Pairs of pixels, Δ*x* = 6 pixels apart, are then correlated over time to calculate the pCF carpet (Fig. 4A, bottom panels). A positive correlation occurs only when a Dl-GFP molecule that is detected in a pixel at position *x* = *x*_1_ and time point *t*_1_ is detected again Δ*x* = 6 pixels to the right at time point *t*_2_. Statistically, all occurrences of positive correlations starting at *x*_1_and having a time difference of Δ*t* = *t*_2_ − *t*_1_ are averaged over 200,000 scans of the line across the nuclei, resulting in a value on the pCF carpet at coordinate (*x*_1_, Δ*t*). To ensure robustness of the data, pCF carpets were also calculated for Δ*x* = 7 or 8 pixels, as well as for pairs of pixels in the right-to-left direction.

**Figure 4:**
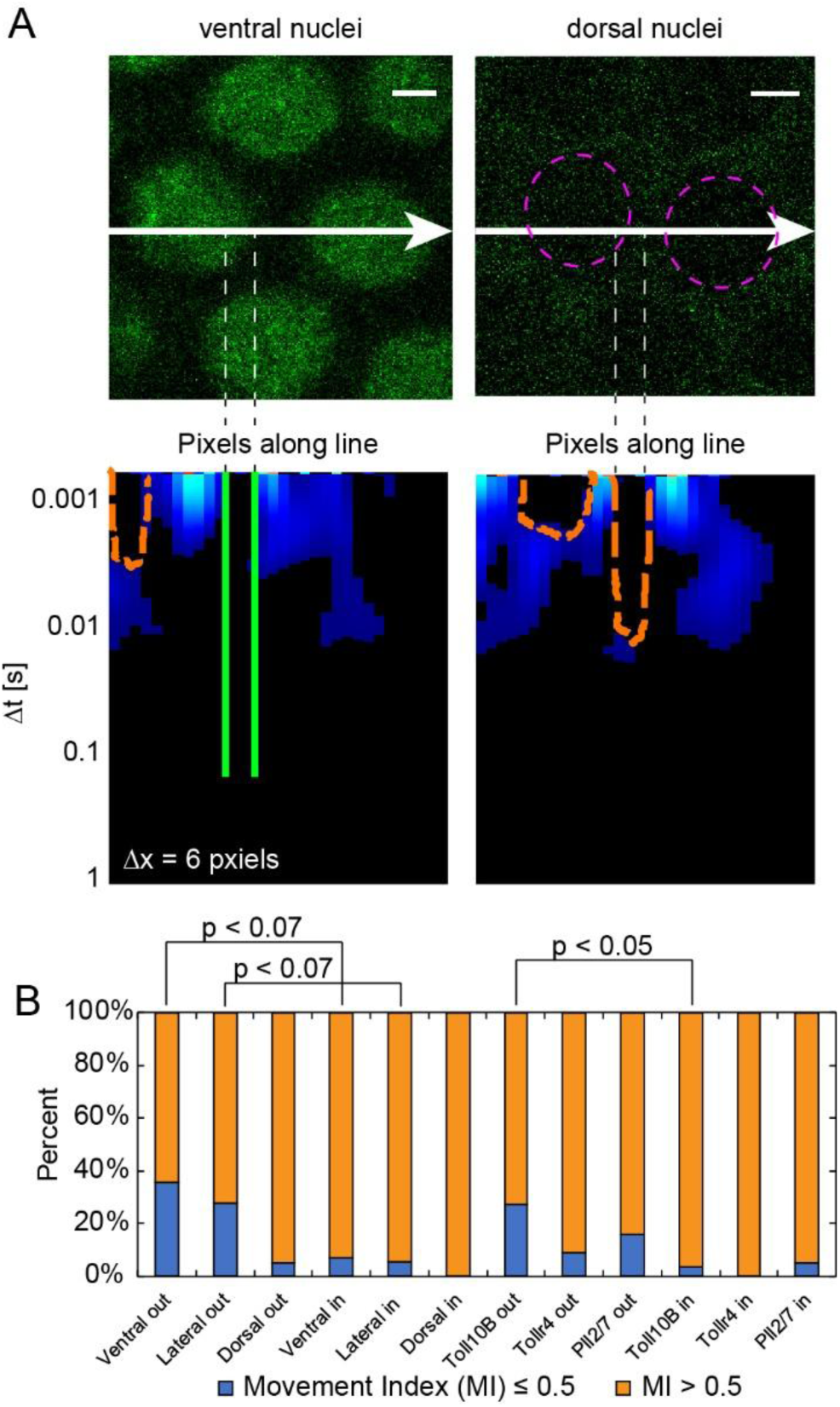
pCF analysis reveals a dorsal-to-ventral asymmetry in nuclear export of Dl. (A) Representative images of Dl-GFP on th ventral (top left image) and dorsal (top right image) sides of the embryo with their respective pCF carpets (bottom left and bottom right). Scale bar = 2um. White arrows represent the direction and y-coordinate of the pCF line scan. Magenta circles in dorsal side image outline the two nuclei crossed by the line scan. Dashed vertical lines illustrate two cases of locations in the images where a crossing of the nuclear envelope is observed (inside the nucleus to outside). A Dl-GFP molecule observed on the left vertical line (inside the nucleus) may or may not be detected at the location of the right vertical line (outside) at a later time point. Orange, dashed lines in the carpet label arches which indicate delayed Dl-GFP movement, while green, solid lines demarcate black regions of the carpet which indicate no Dl-GFP movement. (B) Percent of embryos that have high (Movement Index (MI) > 0.5, orange) and low (MI ≤ 0.5, blue) Dl-GFP movement either out of (out) or into (in) the nucleus. * denotes p<0.07, ** denotes p<0.05, Chi-squared test with likelihood ratio.

After visual inspection of the pCF carpets, we noticed that pixel locations just to the left of the nuclear membrane aligned with either zero correlation (green bars, Fig. 4A), in which no positive correlation is detected over the time course of the experiment, or delayed correlation (orange arches, Fig. 4A), in which the earliest positive correlations are detected much later compared to other pixel locations. These visual inspections suggested that Dl-GFP molecules in the nucleus either failed to cross the nuclear membrane or could not do so immediately. To quantify this, we calculated the Movement Index (MI) for each embryo, which represents the degree of Dl-GFP movement across the nuclear envelope (see Materials and Methods). A high movement index suggests that Dl-GFP can cross the nuclear envelope, which is represented by an arch in the pCF carpet (demarcated by orange, dashed lines in Fig. 4A). In contrast, a low movement index suggests that Dl-GFP movement into or out of the nucleus is blocked, which is represented by a fully black column in the pCF carpet (demarcated by green, solid lines in Fig. 4A). Note that, as pCF tracks the same molecule twice (Digman & Gratton, 2009; Hinde et al., 2010), the MI correlates with the time scale over which an individual Dl-GFP molecule can exit the nucleus, and is not equivalent to a nuclear export rate constant, which is a measure of average nuclear export.

We separated our images into two groups, those with high movement (MI > 0.5) and those with low movement (MI ≤ 0.5). We found that Dl-GFP can move into the nucleus across the spatial regions of the embryo, as the ventral, lateral, and dorsal regions all have a majority of images with high MI values (ventral: 92.86%; lateral: 94.44%; dorsal: 100.00%) (Fig. 4B). However, the ventral and lateral regions have significantly fewer images with high MI values when Dl-GFP is measured leaving the nucleus (ventral: 64.29%; lateral: 72.22%; p<0.07, Chi-squared test with likelihood ratio), suggesting that Dl-GFP movement out of the nucleus is restricted in the ventral and lateral regions of the embryo (Fig. 4B). In contrast, on the dorsal side of the embryo, almost all the images measuring Dl-GFP movement out of the embryo have high MI values (94.44%). To test whether this restricted movement out of the nucleus may be dependent on Toll signaling, we also performed pCF analysis in *Toll^10B^*, *Toll^r4^*, and *pll^2/7^*mutant lines (Fig. S3A), and similarly found that a lower proportion of images in “ventral-like” mutants have high MI values when Dl-GFP movement is measured out of the nucleus (*Toll^10B^*: 96.15% in versus 72.73% out; p<0.05, Chi-squared test with likelihood ratio) (Fig. 4B). This suggests that the restriction of Dl to exit the nucleus correlates with the spatial gradient in Toll signaling. As Dl/DNA interactions also correlate with Toll signaling, DNA binding may explain our pCF results. For example, it is likely that a Dl-GFP molecule detected by pCF in a ventral nucleus may be bound to DNA initially, implying it will take longer on average to exit the nucleus than a Dl molecule not initially bound to DNA. Therefore, on the ventral side, the time scale on which an individual Dl-GFP molecule exits the nucleus is longer than on the dorsal side (where DNA binding is not happening). This can be true even if the nuclear export rate constants are the same on the ventral side vs the dorsal side. Thus, our pCF results are consistent with a high fraction of immobilized Dl in the nuclei on the ventral side of the embryo.

### Analysis of *dl* mutant alleles suggests Cact is present in the nuclei

Our results demonstrate that, on the ventral side, a significant fraction of Dl is bound to DNA, while on the dorsal side, that fraction is statistically indistinguishable from zero. This phenomenon raises the question of how Dl, as a transcription factor, would be present in the nucleus, but not bind to DNA. More generally, we ask what mechanism may cause the fraction of Dl bound to DNA to positively correlate with nuclear Dl concentration, when a mechanism consisting solely of saturating biochemical binding interactions predicts the opposite. As the gradient in Dl/DNA interactions is dictated by Toll signaling, we reasoned that, on the ventral side, Dl/DNA binding may be enhanced by phosphorylation of Dl by Toll. Toll phosphorylation of Dl is known to enhance the affinity of Dl for the nucleus (Fig. 5A’) (Drier et al., 1999), but its effect on DNA binding is unknown. To test this hypothesis, we made another GFP-fused allele of *dl*, this time with an alanine replacing serine at Dl residue 317 (*dl*^S317A^-*gfp*). This allele has been shown to have reduced Toll-mediated phosphorylation of Dl (Drier et al., 1999). Because Dl acts as a homodimer, and thus, like *dl*^R063C^-*gfp*, this allele may be antimorphic (Drier et al., 1999), we used the *nos*-Gal4 line to drive the expression of UAS*-dl*^S317A^-*gfp* (Brand & Perrimon, 1993). As this allele has been shown to drastically reduce the nuclear localization of Dl (Drier et al., 1999), we imaged embryos carrying one copy of this UAS-driven allele and zero copies of wildtype Dl and found that these embryos had no discernible Dl gradient (Fig. S4A). Therefore, to maintain DV polarity so that the ventral and dorsal sides of the embryo could be distinguished, we overexpressed this allele in the background of the BAC recombineered Dl-mGFP construct. We again used the UAS-*dl^wt^-gfp* as a direct wildtype control. As such, if Toll phosphorylation of Dl is responsible for allowing Dl/DNA interactions, then this allele should have reduced DNA binding on the ventral side compared to the wildtype control. However, upon performing ccRICS analysis, we found that the DNA binding of this allele was not statistically distinguishable from the control on the ventral side (Fig. 5B). We note that we cannot completely rule out the hypothesis that Toll signaling has a small effect on Dl/DNA binding, given these embryos expressed the *dl*^S317A^-*gfp* allele in the background of the BAC recombineered construct.

**Figure 5:**
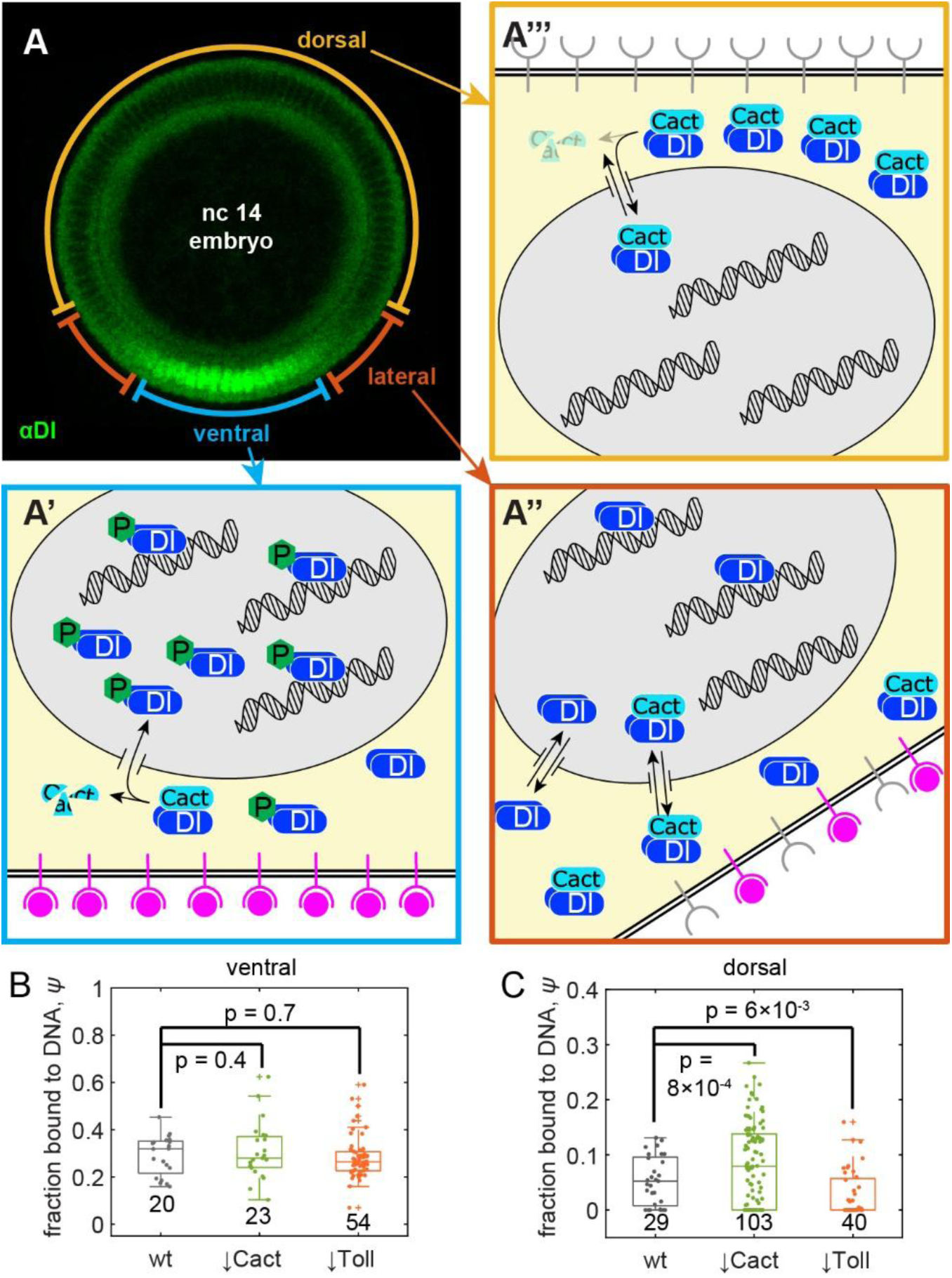
Variation of Dl mobility along the DV axis is due to presence of Cact in the nuclei. (A) (top left) Depiction of the different embryo domains, ventral (blue region), lateral (orange region), and dorsal (yellow region). Same embryo as in Fig. 1A. (A’) Schematic of ventral side of embryo. High levels of activated Toll (magenta receptors) result in the destruction of Cact and phosphorylation of Dl. Free Dl enters the nucleus, so GFP fluorescence is mostly nuclear. Free Dl in the nucleus binds to DNA, which lowers the overall mobility of Dl species, and exits the nucleus at a slow rate. (A’’) Schematic of lateral domain of embryo. Lower levels of Toll activation lead to a mixture of free and Cact-bound Dl. Nuclear and cytoplasmic levels of GFP fluorescence are roughly equal. Dl has a higher mobility due to lower fraction of Dl bound to the DNA. Dl exits the nucleus faster than on the ventral side. (A’’’) Schematic of the dorsal domain of the embryo. Little-to-no free Dl is present, so Dl-GFP fluorescence is mostly cytoplasmic. High mobilities stem from no DNA binding. (B,C) Boxplots of ψ, the fraction of Dl bound to DNA for the UAS-*dl^wt^-gfp* (control), UAS-*dl*^S234P^-*gfp* (reduced Cact binding: ↓Cact), and UAS*-dl*^S317A^-*gfp* (reduced Toll phosphorylation: ↓Toll) alleles on the ventral (B) or dorsal (C) sides. The median line for S317A in (C) is located at ψ = 0. Solid dots represent individual measurements. *p*-values for pairwise comparisons to the control are indicated on the graph. Sample sizes indicated on graph.

Alternatively, we reasoned that, on the dorsal side, Dl in the nuclei may be in complex with Cact, preventing it from binding to DNA (Fig. 5A’’’), as suggested by our experimental and modeling work (Al Asafen et al., 2020; O’Connell & Reeves, 2015; Schloop et al., 2025). As Dl/Cact interactions are downstream of Toll signaling, this hypothesis would be consistent with the gradient in Dl/DNA interactions being dictated by Toll signaling. Therefore, we made a GFP-fused *dl* allele that has reduced Cact binding by mutating serine 234 to a proline (*dl*^S234P^-*gfp*)(Drier et al., 2000) and as above, to avoid potential antimorphic effects, we used *nos-Gal4* to overexpress the allele. It has been reported that eliminating the Dl/Cact binding results in a weak gradient of Dl, with Dl significantly entering the nuclei on the Dl side of the embryo (Drier et al., 2000). As such, we imaged embryos carrying one copy of UAS-*dl*^S234P^-*gfp* without the benefit of the background of the BAC recombineered construct, and found no discernible Dl gradient, similar to the Toll phosphorylation mutant (Fig. S4B). Therefore, to ensure DV polarity of the Dl gradient is maintained, we performed ccRICS on embryos overexpressing this allele in the background of the BAC recombineered Dl-mGFP construct. If the presence of Cact in the dorsal nuclei is responsible for preventing Dl/DNA interactions, then this allele should have increased DNA binding on the dorsal side compared to the wildtype control. Indeed, our ccRICS analysis found that the DNA binding of this allele was greater than the control on the dorsal side (Fig. 5C). The small effect size was likely due to the presence of a wildtype Dl protein in the form of the BAC recombineered Dl-mGFP construct. Indeed, the small effect size may have prevented us from discerning a statistically significant effect of this allele to increase Dl binding on the ventral side (Fig. 5B). Nevertheless, the effect on the dorsal side (Fig. 5C) was statistically significant (p-value ∼ 10^-3^). We conclude that the gradient in nuclear Dl mobility is due to Dl/Cact complex in the nucleus; however, we cannot rule out an additional, albeit small, role that Toll phosphorylation may also play. Our previous modeling work has predicted that the presence of Cact in the nucleus affects the spatial range of the Dl gradient and the robustness of gene expression (Al Asafen et al., 2020; O’Connell & Reeves, 2015). Thus, our findings affect the foundation of our understanding of Dl-dependent patterning of the DV axis.

## Discussion

In this study, we used fluorescence fluctuation spectroscopy methods to quantify the movement of a BAC-recombineered, GFP-tagged Dl protein in the 1-3 hr old *Drosophila* embryo (Fig. 5A). Our investigations ultimately unveiled a DV asymmetry in the mobility of Dl, which was correlated to a similar DV asymmetry in the ability of Dl to bind to DNA. First, through RICS analysis, we identified variations in Dl’s apparent diffusivity along the DV axis within the nucleus. Given the possibility that at least two pools of Dl exist in the nucleus (free vs immobilized), we built a two-component model that accounts for both pools of Dl and found that the DV asymmetry in the fraction of immobilized Dl explained the DV asymmetry in apparent diffusivity. Therefore, to test whether the observed immobilized Dl could be explained by Dl/DNA binding, we used ccRICS to cross-correlate Dl-GFP with Histone-RFP. We indeed found the DV variations in the fraction of immobilized Dl correlated with Dl/DNA binding, with the highest DNA binding on the ventral side. Moreover, the results obtained from pCF showed that there is also variation in the movement of Dl out of the nuclei: on the ventral side, the propensity of Dl to exit the nucleus is not as rapid as it is on the dorsal side. Taken together, our results suggest that there is a DV asymmetry in the efficiency with which Dl can bind to the DNA, and that the restricted nuclear Dl movement on the ventral side of the embryo is explained by a high proportion of DNA-bound Dl.

We identified two plausible explanations that could be causing the DV variation in Dl/DNA interactions. First, Toll-mediated phosphorylation of Dl on the ventral side of the embryo might increase its DNA-binding affinity. Indeed, phosphorylated Dl has increased ability to enter the nucleus (Drier et al., 1999, 2000; Gillespie & Wasserman, 1994; Whalen & Steward, 1993). However, no previous evidence has been shown supporting phosphorylated Dl having an increased affinity for DNA. Second, it is possible that nuclear Dl on the dorsal side is predominantly in complex with Cact, which would result in high average mobility, owing to the fact that it cannot bind DNA (Geisler et al., 1992; Kidd, 1992). In support of this explanation, our recent live imaging of an endogenously-tagged Cact showed that, in *Drosophila* embryos, Cact is present in the nucleus (Schloop et al., 2025). In comparison, mammalian IκB, the homolog of Cact, enters the nucleus to regulate NF-κB dimers, which are then rapidly exported from the nuclei (Arenzana-Seisdedos et al., 1995, 1997; Banerjee & Sen, 2006; Carlotti et al., 2000; Hall et al., 2006; Lisiero et al., 2020; Turpin et al., 1999).

To test these competing hypotheses, we generated two mutant alleles of *dl*, one of which reduced phosphorylation of Dl by Toll (Drier et al., 1999), and the other reduced Dl/Cact interactions (Drier et al., 2000). Using ccRICS, we found that, while the reduced Toll allele exhibited little-to-no change in DNA affinity on the ventral side, the reduced Cact allele did indeed have increased DNA affinity on the dorsal side, consistent with the prediction of the second hypothesis. However, to maintain the DV polarity of the embryo, we expressed these mutant alleles in the background of the BAC recombineered Dl-mGFP construct, such that the effects of these alleles were diluted by the presence of a GFP-tagged wildtype Dl protein. Therefore, while can safely reject the null hypothesis that Cact has no effect on DNA binding, it is possible that our failure to reject the null hypothesis that Toll phosphorylation has no effect on the DNA binding of Dl on the ventral side may be due to this dilution effect.

Genetically, the bulk of the embryos in this study maternally expressed one copy of endogenous Dl and one copy of the BAC-recombineered construct, which presents two caveats. First, it is known that tagging Dl with a fluorescent protein alters its gradient (Reeves et al., 2012), which may in part be due to a lowered diffusivity (Carrell et al., 2017). As such, any raw diffusivity reported here may measure as slightly lower than that of untagged Dl. We do not expect the fraction immobile or the fraction bound to DNA to be affected. A second caveat is that the presence of endogenous Dl may increase the variance of our data. As Dl exists as a dimer (Govind et al., 1992; Isoda & Nüsslein-Volhard, 1994), our measurements are likely to reflect an ensemble average of a tagged/untagged heterodimer mixed with a tagged homodimer at a 2:1 ratio. We do not expect the existence of a mixed population of Dl dimers to alter our main conclusions.

It is thought that the primary role for Cact is to tether Dl to the cytoplasm and prevent nuclear translocation. Indeed, loss of Cact function permits intermediate levels of Dl to enter the nuclei, even on the dorsal side of the embryo (Bergmann et al., 1996; Cardoso et al., 2017; Roth et al., 1991). However, recent studies have discovered multiple functions for Cact, including positive roles in the formation of the Dl gradient. For example, Cact degradation on the ventral side of the embryo creates a driving force for a ventrally-directed net flux of Dl/Cact complex, which has the overall effect of shuttling Dl to the ventral side (Carrell et al., 2017). Furthermore, recent experimental and modeling work has suggested that Cact potentiates nuclear translocation on the ventral side of the embryo (Barros et al., 2021; Cardoso et al., 2017).

In addition to these roles for Cact in establishing the Dl gradient, our experimental results in this study suggest that, on the dorsal side of the embryo, Cact is present inside the nuclei and plays a role in preventing Dl from binding DNA. While this specific finding relates to the dorsal half of the embryo, where Dl levels are low and are expected to have minimal activity in regulating gene expression, knowing whether Dl is in complex with Cact in the dorsal-most nuclei is crucial to our understanding of the Dl gradient. Indeed, our previous fluorescence measurements have revealed a gradient that decays to non-zero basal levels on the dorsal half of the embryo (Liberman et al., 2009; Reeves et al., 2012); such a gradient would have a very limited dynamic range, varying by less than one order of magnitude (O’Connell & Reeves, 2015). In light of our results in this paper, one solution to this problem is to assume all of the Dl on the dorsal side of the embryo is Dl/Cact complex, meaning that the gradient of free Dl would have an unknown overall shape, but would decay to near-zero levels. We have shown that such a free Dl gradient would not only have extended spatial and dynamic ranges, but also superior precision and robustness of gene expression compared to the gradient measured by fluorescence (Al Asafen et al., 2020; Liberman et al., 2009; O’Connell & Reeves, 2015; Reeves et al., 2012). In support of the assumption that all Dl is bound to Cact on the dorsal side, we have recently shown that Cact exhibits dynamics similar to Dl on the dorsal side of the embryo, suggesting that, in both cases, it is predominantly Dl/Cact complex that is being measured (Hiremath et al., 2024; Reeves et al., 2012; Schloop et al., 2025).

Furthermore, our recent experimental work showed that Cact is present in all nuclei and is spatially uniform along the DV axis of the embryo (Schloop et al., 2025). As such, our results in this paper can likely be extended to Dl/Cact interactions in all nuclei, implying that measurements of the Dl gradient based solely on nuclear fluorescence levels are in fact a combination of a spatially graded pool of free (active) Dl and a spatially uniform level of Dl/Cact complex along the entire DV axis. Thus, the “true” Dl activity gradient remains unquantified, and must be determined by deconvolving Dl/Cact complex from free Dl (O’Connell & Reeves, 2015). If all (or nearly all) of the Dl in the dorsal-most nuclei is indeed bound by Cact, the gradient of free Dl, along the entire DV axis, could be recovered by simply subtracting off the basal levels. However, it is likely that some fraction of Dl in the dorsal-most nuclei is free. Thus, to fully understand the positional information carried by the Dl gradient, direct quantification of Dl/Cact complex, perhaps through ccRICS measurements on embryos in which both Dl and Cact are labeled, should be the next step towards gaining future insights into the operation of the highly conserved NF-κB/Dl module. Ultimately, our work highlights the need for quantitative measurements of biophysical parameters to further our understanding of signaling and gene expression, and calls for similar studies in other signaling pathways in embryonic patterning.

## Materials and Methods

### Fly Stocks and Preparation

*Drosophila melanogaster* stocks were kept on corn meal molasses food in vials or bottles at 25°C and all crosses were performed at 25°C. *Drosophila* embryos, that were about 1-2 hrs old, were collected and mounted on slides for imaging. Briefly, flies were left to deposit eggs on fresh grape juice agar plates with yeast paste for 30-60 min. Those embryos were then aged further for 30 min at 25°C to reach the desired developmental stage. Those embryos are then brushed from the grape juice agar plates into a mesh basket using a paintbrush and DI water. The embryos were dechorionated using bleach and then washed with DI water (Carrell & Reeves 2009).

*dl-mGFP* (Dl-GFP) was created by BAC recombineering and *dl*^1.2.5^ flies were generated by cleaning up *dl*^1^ via two homologous recombinations with wildtype flies (Carrell et al. 2017). Flies carrying *dl*^1.2.5^ were crossed to flies carrying H2Av-RFP on the second chromosome (w[*]; P{w[+mC]=His2Av-mRFP1}II.2; BS# 23651) to generate flies that have *dl*^1.2.5^,H2Av-RFP transgene on the second chromosome. H2Av-RFP,*dl*^1.2.5^*,dl-mGFP* flies were created by homologous recombination (Carrell et al. 2017). Thus, the “Dl-GFP embryos” analyzed in this study had a maternal genotype of H2Av-RFP,*dl*^1.2.5^*,dl-mGFP*/*CyO*: one copy of endogenous *dl* and one copy of *dl-mGFP*. The generation of *Toll^r4^* and *Toll^10B^* mutant embryos has been previously described (Stathopoulos et al. 2002). Other *Drosophila* strains were obtained from the Bloomington (BL) stock center, namely *pll^2^* (e[1] pll[2] ca[1]/TM3, Sb[1] BS# 3111) and *pll^7^* (e[1] pll[7] ca[1]/TM3, Sb[1]; BS# 3112), and nos-Gal4 (w[1118]; P{w[+mC]=GAL4::VP16-nanos.UTR}CG6325[MVD1]; BS# 4937).

The pUAST plasmids containing *dl* alleles (*dl*^wt^-*mgfp*, *dl*^R063C^-*mgfp*, *dl*^S317A^-*mgfp*, and *dl*^S234P^-*mgfp*) were prepared by site-directed mutagenesis using appropriate primers with the Q5^®^ Site-Directed Mutagenesis Kit (New England Biolabs Inc., Frankfurt, Germany). All resulting plasmids were confirmed by direct sequencing. The Gal4/UAS overexpression embryos were maternally H2Av-RFP,*dl*^1.2.5^*,dl-mGFP*/*dl*^4^; *nos*-Gal4/UAS-*dl^x^-mGFP*, where *x* is either the R063C, S317A, or S234P mutation, or wildtype. Thus, these embryos maternally carried zero endogenous copies of *dl*, one BAC-recombineered *dl-mGFP* construct, and one UAS-*dl^x^-mGFP* construct.

### Fly Transformation

*Drosophila* embryos were injected with 0.5 ng/µl of each pUAST construct using the PhiC31 method by GenetiVision (Texas, USA). Transformed flies include insertion on chromosome III. Stocks were established for each of these flies. The flies were then crossed into a dl^4^ heterozygous background (dl[4] pr[1] cn[1] wx[wxt] bw[1]/CyO BS# 7096) and finally crossed with dl1.2.5 flies (Carrell et al., 2017) driven by nos-gal4 (BS# 4937, see above). The females from this cross were kept in a cage and the resulting embryos were used for live imaging.

### Mounting and Imaging of *Drosophila* Embryos

*Drosophila* embryos that were 1-2 hrs old were mounted laterally on a microscope slide using a mixture of heptane glue (Supatto et al. 2009) and two pieces of double-sided tape to prevent sample movement (Carrell & Reeves 2015). The embryos were mounted in random orientations along the DV axis. The DV position of each imaged region was estimated by measuring the Dl nuclear-to-cytoplasmic ratio (NCR) of intensity, which varies predictably with position along the DV axis. We categorized positions with NCRs > 1.25 as ventral, positions with 0.75 < NCR < 1.25 as lateral, and positions with NCR < 0.75 as dorsal (Fig. 1A). Embryos were imaged on either a Zeiss LSM 710 or LSM 900 confocal microscope using a 40x 1.2NA water objective. Embryos in nuclear cycle 12 to 14 were selected using the H2Av-RFP marker. Embryos undergoing cell division (as visualized by the H2Av-RFP marker) were not used in for analysis as cell division alters the endogenous Dl distribution.

For image acquisition consistent with RICS analysis for the data shown in Figs. 1-3, using a Zeiss LSM 710, a 256 x 256 pixel region of interest of the embryo was selected for measurements (Supplemental Figure 1). The pixel size of the region varied from 40 nm to 100 nm to include different numbers of nuclei. The 25 mW 488 nm laser intensity ranged from 0.5% to 3.0% and the 20 mW 570 nm laser intensity ranged from 0.1% to 0.2%. The region of interest was raster scanned with a pixel dwell time of 6.3 μs for 200 frames (total imaging time of approximately 1.5 minutes). For dl-mutants (data shown in Fig. 5) we used Zeiss LSM 900 with a 1024 x 1024 pixel region of interest, with a pixel size of 30 nm (Supplemental Figure 1), having the laser intensity range same as mentioned above for the 10 mW 488 nm laser and the 10 mW 561 nm laser. The region of interest was raster scanned with a pixel dwell time of 2.1 μs for 100 frames (total imaging time of approximately 9 min). The range of imaging parameters is reported in Table S1.

For image acquisition consistent with pCF analysis, a 32×1 line scan through 2 to 4 nuclei was used as the region of interest (Fig. 4A). The pixel size was not set for each image but, instead, varied between 40-100nm according to the region of interest selected. The laser intensities varied as in the RICS imaging. The line was scanned with a pixel dwell time of 6.30µs for 200,000 to 400,000 time points (total imaging time of approximately 1.5 to 3 minutes). The range of imaging parameters is reported in Table S1.

### Raster Image Correlation Spectroscopy (RICS) analysis using MATLAB implementation

RICS analysis of time series images is briefly described here (see Supplemental Information for more details). For each frame *j* (of size 256×256 or 1024×1024 pixels) in the Dl-GFP channel, the scanning ACF, *G*_*j*_(*ξ*, *η*), was computed according to the following for all pixel shifts *ξ*, *η*:

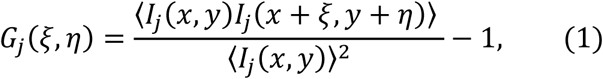

where *I*_*j*_(*x*, *y*) is the intensity of frame *j* at pixel coordinates (*x*, *y*) and the angle brackets denote ensemble average in both *x* and *y*. In practice, *G*_*j*_(*ξ*, *η*) is evaluated using the fast Fourier transform method (Digman & Gratton, 2011). The final ACF for total RICS, *G*_*s*_(*ξ*, *η*), was computed as the average of *G*_*j*_(*ξ*, *η*) for all frames *j* in the time series. Cross-correlation functions were computed in a similar way, except *I*_*j*_(*x* + *ξ*, *y* + *η*) was replaced by the analogous array from the H2Av-RFP channel, and the 〈*I*_*j*_(*x*, *y*)〉^2^in the denominator was replaced by the product of the average intensity in the Dl-GFP channel and the average intensity in the H2Av-RFP channel.

### Fitting models to the ACFs

The one component diffusion model for the RICS ACF is (Fig. 1D-F; Digman et al., 2005b):

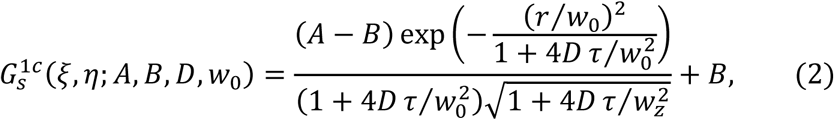

where 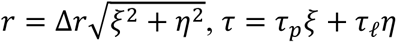, Δ*r* is the pixel size in microns, *τ*_*p*_ and *τ*_ℓ_ are the pixel dwell time and line scan time in seconds, respectively, and *w*_0_and *w*_*z*_are the waist sizes of the point spread function in the planar and axial directions, respectively. The two component diffusion/binding model is a linear combination of two instances of Equation (2):

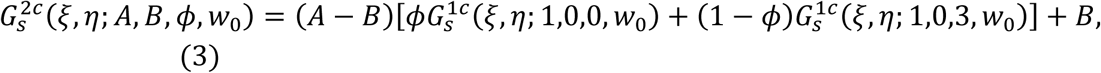

where *ϕ* is the fraction of Dl-GFP that is immobilized, in the first term, *D* is held fixed at zero, and in the second term, *D* is held fixed at 3 μm^2^/s. Within both terms, 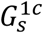 is normalized such that *A* = 1 and *B* = 0, because the *A* and *B* parameters are otherwise present in Equation (3).

Models of the ACF were fit to the ACFs calculated from the total, nuclear, or cytoplasmic data in two steps. First, the fast direction of the ACF (*η* = 0) was used as target to fit 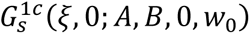, that is, Equation (2) with *η* = 0 and *D* = 0. The shape of this function is purely Gaussian and approximates the point spread function (PSF). This fit is highly robust, and performing this fit first allows for accurate estimation of the parameter *A*. Two parameters, the PSF waist size, *w*_0_, and the background level, *B*, are theoretically fixed; however, we allowed them to vary slightly from their theoretical values of 0.25 μm and zero, respectively, for robustness of fit.

In the second step, with *A* from the first step fixed, either Equation (2), for the one-component model, or Equation (3), for the two component model, was fit to *G*_*s*_(*ξ*, *η*) using least squares and varying *B*, *D*, *w*_0_ or *B*, *ϕ*, *w*_0_, respectively. The parameters Δ*r*, *τ*_*p*_, *τ*_ℓ_, *w*_*z*_ were microscope parameters and considered fixed. More information can be found in the Supplemental Information.

### Pair Correlation Function (pCF) analysis using SimFCS software

Acquisitions were taken of 200,000 scans of a line 32 pixels across (pixel size 0.5 μm), with a pixel dwell time of 6.3 μs and a line time of 0.47 ms, on either the dorsal, lateral, or ventral sides of nc 14 embryos. To ensure minimal movement of nuclei occurred during the line scan aquisition, 512×512 images were taken before and after each acquisition, where the middle y-coordinate of these images corresponded to the location of the line scan (white arrows in Fig. 4A). pCF analysis of the collected line scans was performed using the SimFCS software (Digman et al, 2005; https://www.lfd.uci.edu/globals/) as described in Clark and Sozzani, 2017. Three pixel distances of 6, 7, and 8 pixels were used as technical replicates to account for differences in sizes of nuclei. If Dl-GFP showed movement out of/into a majority of nuclei (no black region or black arch in the pCF carpet), that image was assigned a movement index of 1. Otherwise, if the majority of Dl-GFP showed no movement (black vertical region in the pCF carpet), that image was assigned a movement index of 0 (Fig. 4A). The 3 technical replicates were then averaged for each biological replicate. The analysis was repeated for the line scan in the reverse direction, resulting in 6 total technical replicates per image.

### Quantification and statistical analysis

Non-linear least squares was used to fit Equation (2) to the RICS ACFs. The fitted diffusivity value is *t*-distributed, and a standard error for the diffusivity (radius of a 68% confidence interval) was calculated. With the exception of five time course image stacks of nuclear RICS either on the dorsal side of wildtype embryos, or in *pll*^2/7^ embryos (the two cases in which nuclear intensity is very low), for all fitted diffusivities, the standard error was less than 4 μm^2^/s. This was used as a filter for the data; the five time course image stacks in which the standard error was greater than 4 μm^2^/s were removed from the analysis.

For the diffusivity of Dl calculated by RICS (Fig. 1D-F), the ccRICS data (ratio of amplitudes; Fig. 3C,D), and for the correlation between nuclear diffusivity and φ (Fig. 4E), simple linear least squares regression was applied. The slope of the regression trend line is t-distributed with *n*-2 degrees of freedom, where *n* is the number of data points. Reported *p*-values are for *t*-tests with the null hypothesis that the slope is zero.

For the pCF data (Fig. 2), a Chi-squared test with likelihood ratio was used to test if the proportion of Movement Index (MI) values that were less than 0.5 was different for movement measured into the nucleus vs movement measured out of the nucleus. The Chi-squared test was performed using JMP software (https://www.jmp.com/).

For all tests, a p-value less than 0.05 was considered significant, and for the Chi-Squared test we also consider the two p-values that were less than 0.07 as significant (Fig. 2). The statistical tests performed on each sample are listed in the Results text as well as the figure legends. Sample sizes are indicated in the figures. Means and standard errors are reported in the Results.

## Data and software availability

Images, tables, and sequencing results will be uploaded to a data repository, Data Dryad. Accession numbers will be available upon notification of acceptance for publication. Matlab codes will be uploaded to github and will also be available from the Reeves Lab website.

## Acknowledgements

The authors acknowledge the use of the Cellular and Molecular Imaging Facility (CMIF) at North Carolina State University, which is supported by the State of North Carolina and the National Science Foundation.

