## Supplemental text for "Dorsal/NF-κB exhibits a dorsal-to-ventral mobility gradient in the Drosophila embryo"

### Supplemental Information for "Dorsal/NF-κB exhibits a dorsal-to-ventral mobility gradient in the Drosophila embryo"

##### *Calculating RICS ACFs and ccRICS CCFs from image data*

Time series images consistent with RICS analysis were analyzed in the following way. Let $I_{0,j}$ be the image frame in the Dl-GFP channel at time point $t_j$. A background subtraction by $I_{sw,j}$, the average of a sliding window with a five-frame radius of frame $j$, was performed to remove the effect of global movement of large structures (M. a Digman et al., 2008): $I_{bs,j} = I_{0,j} - I_{sw,j}$. Next, the average intensity of every pixel in the sliding window, $\bar{I}_{sw,j}$, was added back to frame $j$ to ensure the frame intensity was not near zero, to yield array $I_j = I_{0,j} - I_{sw,j} + \bar{I}_{sw,j}$. Next, a 2D fast Fourier transform of the frame was computed to yield the array $J_j = \mathcal{F}[I_j]$ in the Fourier domain. Next, an array $g_j$ was computed as the inverse Fourier transform of $J_j$ multiplied by its complex conjugate on an element-by-element basis: $g_j = \mathcal{F}^{-1}[J_j J_j^*]$. Finally, the autocorrelation function, $G_j$, for frame $j$ was computed as $G_j = g_j/(HW\bar{I}_j^2) - 1$, where $H$ and $W$ are the original height and width of the image frame, and $\bar{I}_j$ is the mean intensity of all pixels in $I_j$. The final autocorrelation function, $G_s$, was computed as the average of all $G_j$ for $j = 1 \ldots n_t$, where $n_t$ is the number of frames in the image acquisition. Cross-correlation functions were computed in a similar way, except $J_j$ was multiplied by the complex conjugate of the Fourier transform of frame $j$ from the background-subtracted H2A-RFP channel.

Due to the fact that photon shot noise adds to the occupation variance at position (0,0), the computed value of the autocorrelation function at position (0,0) is incorrect. Additionally, noise levels can also affect the value of $G_s$ at position (1,0) (i.e., at $\xi = 1, \eta = 0$). For plotting purposes, both of these pixels are interpolated by a quadratic with symmetry at $\xi = 0$ using the points at (2,0), (3,0), and (4,0). This is the same procedure used in the simFCS software (Gratton, personal communication). For model fitting purposes, both of these points are ignored.

##### *Calculating nuclear and cytoplasmic RICS ACFs and nuclear ccRICS CCFs*

To find nuclear RICS, a nuclear mask for each frame $j$ was found in the following manner. First, frames one through $n_t$ were divided into groups of ten frames each. In each group $k$, the ten frames were averaged to produce an average frame (to improve nuclear segmentation accuracy). This average frame of group $k$ was eroded with a disk-shaped structuring element of one micron, dilated with a disk-shaped element of a half micron, then Gaussian blurred with a standard deviation of a half micron, which resulted in a frame called $I_{EDG,k}$.

Next, the frame $I_{EDG,k}$ was saturated by 2% at both ends. In other words, the 2$^{nd}$ and 98$^{th}$ percentile intensities, $i_2$ and $i_{98}$, respectively, were found, and $I_{EDG,j}$ was rescaled by:

$$I_{rs,k} = \frac{I_{EDG,k} - i_2}{i_{98} - i_2}$$

This resulted in a frame with 2% of its pixels less than zero, and 2% greater than one. These were set to zero and one, respectively.

Next, a global hard threshold was chosen based on the microscope zoom-level and time point (to account for possible bleaching of the H2A-RFP channel). If the pixel size was less than 0.05 microns, the threshold to determine the nuclei varied linearly between 0.30 for $k = 1$ and 0.15 for $k = 20$, while the threshold to determine the cytoplasm varied linearly between 0.1 for $k = 1$ and 0.05 for $k = 20$. If the pixel size was greater than 0.05 microns, the global thresholds were time invariant: 0.55 for nucleus, and 0.25 for cytoplasm. After global thresholding of $I_{rs,k}$, a hole-filling operation was performed on the nuclei, resulting in $I_{nuc0,k}$. This mask was then eroded by 0.5 microns to produce the final nuclear mask, $I_{nuc,k}$ for group $k$.

To find the cytoplasmic mask, the binary image $I_{nuc0,k}$ was inverted, and the resulting image was eroded by 0.5 microns, producing the final cytoplasmic mask, $I_{cyt,k}$, for group $k$. The erosion steps on $I_{nuc0,k}$ and its inverse were to create a conservative estimate of nucleus and of cytoplasm. The nuclear or cytoplasmic mask for group $k$ served as the mask for the frames that composed group $k$.

Once $I_{nuc,k}$ was found for the frames in group $k$, all pixels in the array $I_j$ outside the mask were set to zero, yielding $I_{2,j}$. Then array $g_{nuc,j}$ was computed similar to $g_j$, but using $I_{2,j}$ instead of $I_j$. Additionally, an array, $D_k$, analogous to $g_{nuc,j}$, was calculated using $I_{nuc,k}$ instead of $I_{2,j}$. Finally, the autocorrelation function for that frame and mask was calculated as $G_{nuc,j} = g_j/(D_k \bar{I}_{2,j}^2) - 1$, where the division is on a pixel-by-pixel basis, and where $\bar{I}_{2,j}$ is the mean intensity of all pixels in $I_{2,j}$ within mask $I_{nuc,k}$. The final autocorrelation function, $G_{s,nuc}$, was computed as the average of all $G_{nuc,j}$ for $j = 1 \dots n_t$. Cross correlation functions using a nuclear mask were computed in a similar way as described here and above, except $\bar{I}_{2,j}$ is replaced by the geometric mean of the mean intensities of $I_{2,j}$ and the corresponding H2A-RFP image.

To find cytoplasmic RICS, the same procedures are followed, except mask $I_{cyt,k}$ is used instead of $I_{nuc,k}$.

***Fitting models to the RICS ACFs***

The theoretical RICS autocorrelation function for a one-component, diffusive species, is (Brown et al., 2008; M. A. Digman, Brown, et al., 2005; M. A. Digman, Sengupta, et al., 2005):

$$G_s(\xi,\eta) = \frac{1}{\pi^{3/2} w_0^2 w_z \bar{c}} \left( \frac{1}{1 + \frac{4D(\tau_p \xi + \tau_\ell \eta)}{w_0^2}} \right) \sqrt{\frac{1}{1 + \frac{4D(\tau_p \xi + \tau_\ell \eta)}{w_z^2}}} \exp\left( -\frac{(\xi^2 + \eta^2)\Delta r^2}{w_0^2 + 4D(\tau_p \xi + \tau_\ell \eta)} \right). \quad \text{(S1)}$$

In this formula, $\bar{c}$ is the average concentration of the diffusive species, $D$ is its diffusivity, $\tau_p$ and $\tau_\ell$ are the pixel dwell time and line time, respectively, $\Delta r$ is the pixel size, $\xi$ and $\eta$ are the pixel shifts in the fast and slow directions, respectively, and $w_0$ and $w_z$ are measures of the spatial extent of the excitation density in the $xy$ plane (usually taken to be the radius of the first local minimum in the point spread function) and along the axial ($z$) direction, respectively. Note that the prefactor $1/(\pi^{3/2} w_0^2 w_z \bar{c}) = 1/\bar{n}$ (the average number of molecules in the confocal volume) because the confocal volume is given by $\pi^{3/2} w_0^2 w_z$. Therefore, a measurement of $G(0)$ gives an estimate of the average absolute concentration of the fluorophore in solution. In practice, there is a coefficient, $\gamma = \sqrt{2}/4$, in the numerator of the prefactor, which accounts for the uneven illumination across the voxel (Brown et al., 2008).

One of the unique features of RICS is that, because there are two time scales, the pixel dwell time and the line time, which are embedded within the fast and slow scanning directions, respectively, the RICS 2D ACF captures multiple behaviors in solution (M. A. Digman, Sengupta, et al., 2005). In particular, at a pixel shift of zero in the slow scanning direction, Equation S1 becomes (with the factor $\gamma$ included):

$$G_s(\xi,0) = \frac{\gamma}{\pi^{3/2} w_0^2 w_z \bar{c}} \left( \frac{1}{1 + \frac{4D\tau_p \xi}{w_0^2}} \right) \sqrt{\frac{1}{1 + \frac{4D\tau_p \xi}{w_z^2}}} \exp\left( -\frac{\xi^2 \Delta r^2}{w_0^2 + 4D\tau_p \xi} \right), \quad \text{(S2)}$$

and if we define the diffusion time, $\tau_D = w_0^2/(4D)$, which is the average time for a particle to diffuse a distance of the PSF xy waist, then this equation becomes:

$$G_s(\xi,0) = \frac{\gamma}{\pi^{3/2} w_0^2 w_z \bar{c}} \left( \frac{1}{1+\frac{\tau_p \xi}{\tau_D}} \right) \sqrt{\frac{1}{1+\frac{\tau_p \xi}{\omega^2 \tau_D}}} \exp\left( -\frac{(\xi \Delta r / w_0)^2}{1+\frac{\tau_p \xi}{\tau_D}} \right), \qquad \text{(S3)}$$

where $\omega = w_z/w_0 \sim 3$ . As $D \sim$ 1-10 µm$^2$/s, $w_0 \sim 0.25$ µm, and $\tau_p \sim 5 \times 10^{-6}$ s, then $\tau_p/\tau_D \sim 10^{-3}$ or less. As such, by far the largest effect of the pixel shift in the fast direction, $\xi$, is in the numerator of the Gaussian shape:

$$G_s(\xi,0) = \frac{\gamma}{\pi^{3/2} w_0^2 w_z \bar{c}} \exp(-(\xi \Delta r / w_0)^2), \qquad \text{(S4)}$$

which is just the approximation of the PSF (Elson & Magde, 1974; Krichevsky & Bonnet, 2002). Therefore, we can expect the fast direction of the RICS ACF to closely resemble a Gaussian shape. We used this information to fit our ACF models to the ACF data in a two-step fashion. In the first step, we fit the fast direction to a Gaussian, which gave us a reliable estimate of ACF amplitude. In the second step, we fit the entire ACF to the one- or two-component model, holding fixed the amplitude found in the first step.

In practice, the fast direction is fit to:

$$G_s(\xi,0) = (A-B)\exp(-(\xi \Delta r / w_0)^2) + B, \qquad \text{(S5)}$$

where $A$ is the amplitude, the background parameter $B$ is allowed to vary near zero (between -1e-2 and 1e-2) for robustness of fit, the PSF waist $w_0$ is allowed to deviate from the measured value of 0.25 to increase goodness of fit, and $\Delta r$ is held fixed as reported by the microscope software.

As mentioned in a previous section, the points G(0,0) and G(1,0) are ignored in model fitting. In the slow direction, which is largely controlled by diffusion, small errors in the ACF amplitude can lead to large errors in estimates of diffusivity (for the one-component model) or of the immobile fraction (for the two-component model). Therefore, the first step in the two-step fitting is crucial so that a good estimate of the ACF amplitude can be obtained.

For the second step, the entire ACF is fit to either a one-component model or a two-component model. For the one-component model, Equation S1 is modified to be

$$G_s(\xi,\eta) = (A-B) \left( \frac{1}{1+\frac{4D(\tau_p \xi + \tau_\ell \eta)}{w_0^2}} \right) \sqrt{\frac{1}{1+\frac{4D(\tau_p \xi + \tau_\ell \eta)}{(3w_0^2)}}} \exp\left( -\frac{(\xi^2+\eta^2)\Delta r^2}{w_0^2 + 4D(\tau_p \xi + \tau_\ell \eta)} \right) + B, \qquad \text{(S6)}$$

where $B$ and $D$ are adjustable parameters and where $A$ and $w_0$ are obtained from the first fitting step. We allowed $B$ to vary between -1e-2 and 1e-2 for robustness of fit, and for $D$ to vary between 0 and 30 µm$^2$/s.

For the two component model, the ACF is:

$$G_s(\xi,\eta) = (A-B)\left( \phi \hat{G}_1(\xi,\eta) + (1-\phi)\hat{G}_2(\xi,\eta) \right) + B, \qquad \text{(S7)}$$

where $\hat{G}_i(\xi,\eta)$ is given by Equation S6, normalized such that $A = 1$ and $B = 0$, because the $A$ and $B$ parameters are already present in Equation S7. To perform the fit of S7 to the data, we set $w_z = 3w_0$, and we fixed the diffusivity for $\hat{G}_1$ to be $D_1 = 3$ µm²/s and for $\hat{G}_2$ to be $D_2 = 0$. Again, $B$ was allowed to vary between -1e-2 and 1e-2 for robustness of fit.

In fitting a model to the red channel ACF, in the second step, Equation S6 was used with $D = 0$.

***The cross correlation function from cross-correlation RICS (ccRICS)***

In fluorescence cross-correlation spectroscopy (FCCS) and in cross-correlation RICS (ccIRCS) the fluorescence in one color channel is cross-correlated with the fluorescence in a different color channel. To calculate the cross-correlation function (CCF), it is important to note that the two lasers have different PSF centers and sizes. In this paper, the green fluorescence comes from Dl-GFP and the red fluorescence comes from H2A-RFP. We define $w_{0,g}$ and $w_{z,g}$ be the dimensions of the PSF in the green channel, and $w_{0,r}$ and $w_{z,r}$ be the dimensions of the PSF in the red channel. We assume there are three species in the nucleus: species 1 = free Dl-GFP, species 2 = Dl-GFP bound to DNA, and species 3 = "free" H2A-RFP (that is, H2A-RFP not near DNA-bound Dl-GFP). The theoretical CCF for an immobile species (such as Dl bound to DNA) is (Weidemann et al., 2002):

$$G_{cc,s}(\xi,\eta) = \frac{\bar{c}_2}{\pi^{3/2}\hat{w}_0^2\hat{w}_z(\bar{c}_1+\bar{c}_2)(\bar{c}_2+\bar{c}_3)} \exp\left(-\frac{(\xi\Delta r - d_x)^2 + (\eta\Delta r - d_y)^2}{\hat{w}_0^2} - \frac{d_z^2}{\hat{w}_z^2}\right), \qquad \text{(S8)}$$

In this equation, $\bar{c}_i$ denotes the average concentrations of species $i = 1,2,3$; $d_x, d_y$ are the displacements between the centers of the PSFs of the two lasers in the x,y plane and $d_z$ represents the axial displacement; and the average PSF sizes are defined as $\hat{w}_0^2 = 0.5(w_{0,g}^2 + w_{0,r}^2)$ and $\hat{w}_z^2 = 0.5(w_{z,g}^2 + w_{z,r}^2)$.

When fitting to the CCF data, Equation S8 was modified to be

$$G_{cc,s}(\xi,\eta) = (A_{cc} - B)\exp\left(-\frac{(\xi\Delta r - d_x)^2 + (\eta\Delta r - d_y)^2}{\hat{w}_0^2}\right) + B, \qquad \text{(S9)}$$

where again $B$ was allowed to vary between -1e-2 and 1e-2 for robustness of fit. The model was fit directly, and not by a two-step procedure. This is because, for cross-correlation, $G_{cc,s}(0,0)$ and $G_{cc,s}(1,0)$ are not plagued by shot noise. Note that the $d_z^2/\hat{w}_z^2$ term was ignored because it is predicted to be only a small correction to the true amplitude.

Note that the amplitude of the CCF, $A_{cc}$, is defined as:

$$A_{cc} = \frac{\bar{c}_2}{\pi^{3/2}\hat{w}_0^2\hat{w}_z(\bar{c}_1+\bar{c}_2)(\bar{c}_2+\bar{c}_3)}. \qquad \text{(S10)}$$

Combining this result with the amplitude of the ACF in the red channel, $A_{red}$, we obtain:

$$\frac{A_{cc}}{A_{red}} = \frac{w_{0,r}^2 w_{z,r}}{\hat{w}_0^2\hat{w}_z}\frac{\bar{c}_2}{\bar{c}_1+\bar{c}_2} = \frac{w_{0,r}^2 w_{z,r}}{\hat{w}_0^2\hat{w}_z}\psi. \qquad \text{(S11)}$$

Therefore, from measurements of the amplitudes of the CCF, the red ACF, and PSF sizes, we can calculate $\psi$, the fraction of Dl-GFP bound to DNA.

***Supplemental Figures***

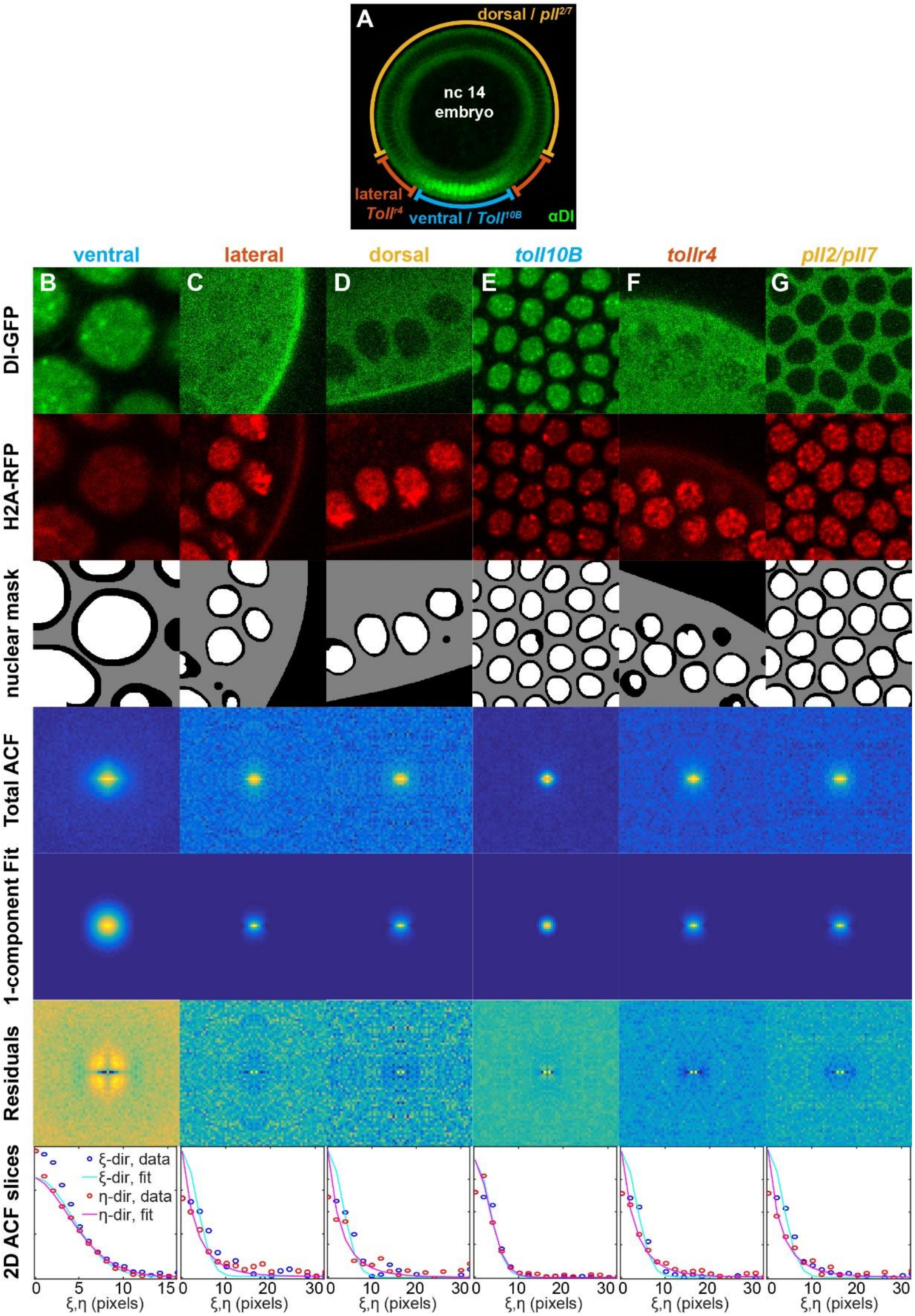

**Figure S1: Representative RICS images and ACFs.** (A) Fixed nc 14 embryo stained for the Dl gradient (same illustration as in Fig. 5). The annotations indicate regions of the embryo that correspond to ventral, lateral, and dorsal. The mutant genotypes included denote that Toll10b is “ventral-like”, Tollr4 is “lateral-like”, and pll2/7 is “dorsal-like”. (B-D) Representative RICS analyses, including snapshot of the Dl-GFP channel and H2A-RFP channel, the nuclear and cytoplasmic masks, the heatmap of the ACF for total RICS, the ACF for the one-component model fit, the residuals to the fit, and the slices of the ACF in the fast and slow directions. Note that the nuclear mask is in white, and the cytoplasmic mask in gray. The residuals are defined as the Total ACF minus the fitted ACF. In the 2D slices plots, open circles are data, and solid curves are the best fit. Blue open circles and cyan solid curve correspond to the fast direction ($\xi$), while red open circles and magenta solid curve correspond to the slow direction ($\eta$). (B) Ventral side. (C) Lateral region. (D) Dorsal side. (E) Toll10b embryo. (F) Tollr4 embryo. (G) pll2/7 embryo. Related to Fig. 1.

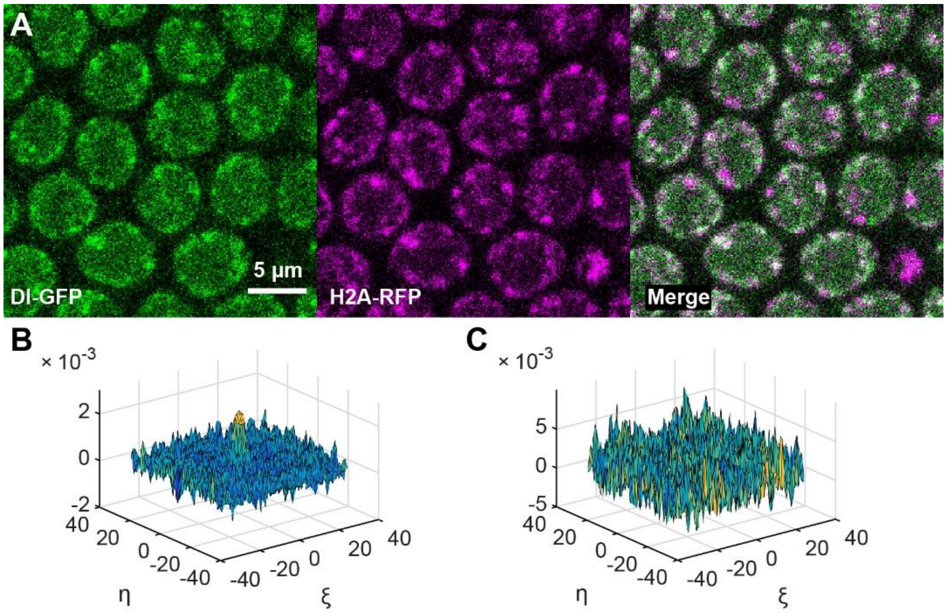

**Figure S2: Cross correlation between Dl-GFP and H2A-RFP.** (A) Snapshot of $Toll^{10b}$ embryo with high correlation between Dl-GFP (green) and H2A-RFP (magenta). The correlation can be seen visually, as noted by the white regions in the nuclei in the merged image. (B) Cross-correlation function for the timecourse shown in (A). (C) Cross correlation function for a timecourse from a $pll^{2/7}$ embryo, in which no correlation is found. Related to Fig. 3.

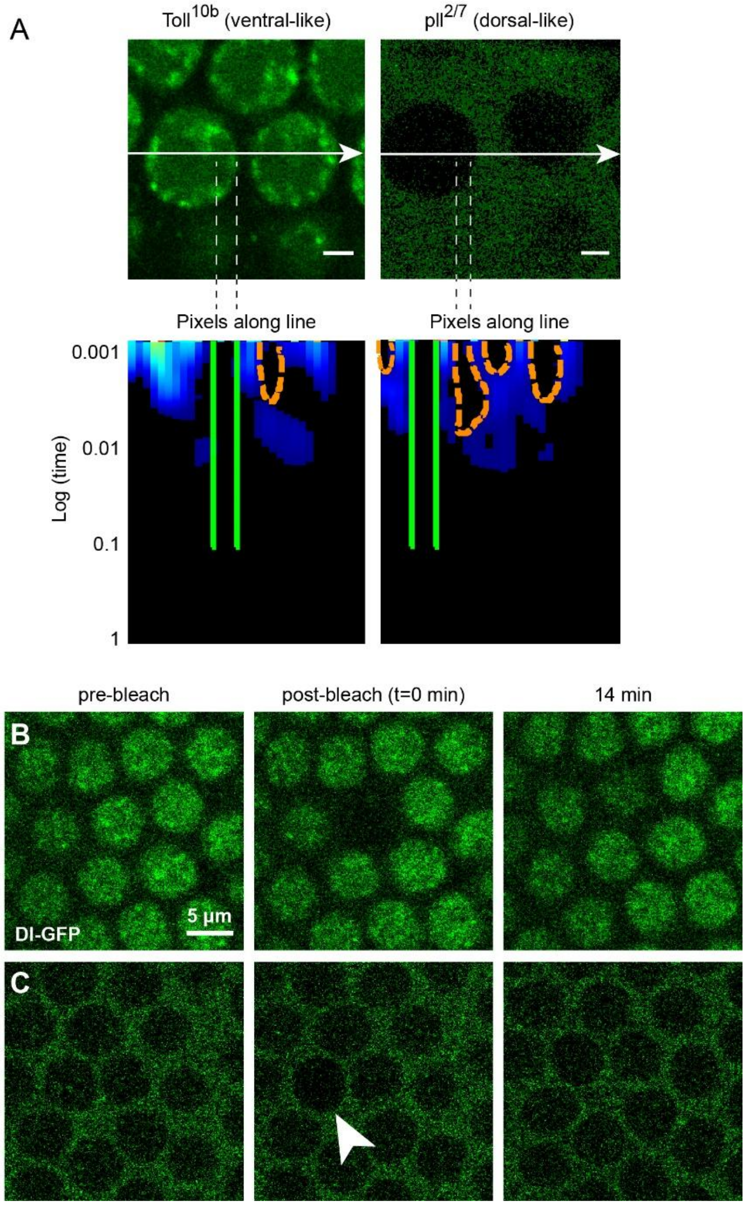

**Figure S3: pCF and FRAP measurements of movement of Dl-GFP in and out of the nucleus.** (A) Representative pCF analyses for *Toll*$^{10b}$ (left column) and *pll*$^{2/7}$ (right column) embryos. Top row: image frame of region where pCF was performed. Bottom row: pCF carpets that correspond to the images in the top row. Scale bar = 2μm. Arrow represents the direction of the line scan. Orange, dashed lines in the carpet label arches which indicate delayed Dl-GFP movement, while green, solid lines demarcate black regions of the carpet which indicate no Dl-GFP movement. Toll10b is "ventral-like" and pll2/7 is "dorsal-like". (B-C) Representative snapshots of FRAP experiments for times given at the top of the images for the ventral (B) and dorsal (C) sides of the embryo. Movies of full timecourses for the embryos in (B) and (C) are in Movies S4 and S5, respectively. Arrowhead in (C) indicates the bleached nucleus.

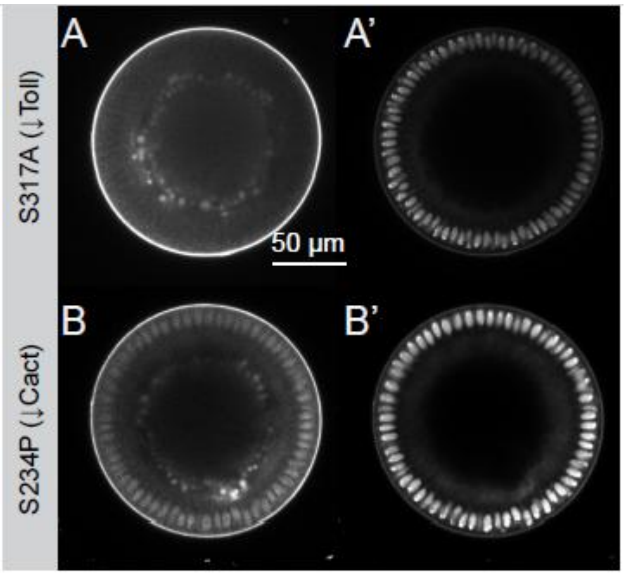

**Figure S4: Dl localization and gradient in embryos with different mutations.** (A, A') The S317A mutation (impairing Toll signaling) shows no discernible Dl gradient. In particular, there are no nuclei that appear "ventral-like", which prevents us from testing the Toll-signaling hypothesis in these embryos. (B, B') The S234P mutation (reducing Cactus binding) demonstrates high levels of Dl in all nuclei, and thus, no nuclei appear "dorsal-like," which prevents us from testing the "Cactus-binding" hypothesis in these embryos.

***Supplemental Movie Captions***

**Movie S1. RICS time course acquisition on the ventral side of a wildtype embryo.** Green: Dl-GFP. Magenta: H2A-RFP. Image frames 26.6 µm on a side.

**Movie S2. RICS time course acquisition on the lateral side of a wildtype embryo.** Green: Dl-GFP. Magenta: H2A-RFP. Image frames 26.6 µm on a side.

**Movie S3. RICS time course acquisition on the dorsal side of a wildtype embryo.** Green: Dl-GFP. Magenta: H2A-RFP. Image frames 26.6 µm on a side.

**Movie S4. RICS time course acquisition of a *$Toll^{10b}$* embryo.** This is an example of a time course with a high amplitude of the cross-correlation function. Green: Dl-GFP. Magenta: H2A-RFP. Image frames 26.6 µm on a side.

***Supplemental References***

Brown, C. M., Dalal, R. B., Hebert, B., Digman, M. A., Horwitz, A. R., & Gratton, E. (2008). Raster image correlation spectroscopy (RICS) for measuring fast protein dynamics and concentrations with a commercial laser scanning confocal microscope. *J Microsc*, *229*(Pt 1), 78–91. https://doi.org/10.1111/j.1365-2818.2007.01871.x

Digman, M. A., Brown, C. M., Sengupta, P., Wiseman, P. W., Horwitz, A. R., & Gratton, E. (2005). Measuring fast dynamics in solutions and cells with a laser scanning microscope. *Biophys J*, *89*(2), 1317–1327. https://doi.org/10.1529/biophysj.105.062836

Digman, M. A., Sengupta, P., Wiseman, P. W., Brown, C. M., Horwitz, A. R., & Gratton, E. (2005). Fluctuation correlation spectroscopy with a laser-scanning microscope: exploiting the hidden time structure. *Biophys J*, *88*(5), L33--L36. https://doi.org/10.1529/biophysj.105.061788

Digman, M. a, Dalal, R., Horwitz, A. F., & Gratton, E. (2008). Mapping the number of molecules and brightness in the laser scanning microscope. *Biophysical Journal*, *94*(6), 2320–2332. https://doi.org/10.1529/biophysj.107.114645

Elson, E. L., & Magde, D. (1974). Fluorescence correlation spectroscopy. I. Conceptual basis and theory. *Biopolymers*, *13*(1), 1–27. https://doi.org/10.1002/bip.1974.360130102

Krichevsky, O., & Bonnet, G. (2002). Fluorescence correlation spectroscopy: the technique and its applications. *Reports on Progress in Physics*, *65*(2), 251–297. https://doi.org/10.1088/0034-4885/65/2/203

Weidemann, T., Wachsmuth, M., Tewes, M., Rippe, K., & Langowski, J. (2002). Analysis of Ligand Binding by Two-Colour Fluorescence Cross-Correlation Spectroscopy. *Single Molecules*, *3*(1), 49–61. https://doi.org/10.1002/1438-5171(200204)3:1<49::AID-SIMO49>3.0.CO;2-T
